# The innate immune sensor Toll-like receptor 2 controls the senescence-associated secretory phenotype

**DOI:** 10.1101/466755

**Authors:** Priya Hari, Fraser R. Millar, Nuria Tarrats, Jodie Birch, Curtis J. Rink, Irene Fernández-Duran, Morwenna Muir, Andrew J. Finch, Valerie G. Brunton, João F. Passos, Jennifer P. Morton, Luke Boulter, Juan Carlos Acosta

## Abstract

Cellular senescence is a stress response program characterised by a robust cell cycle arrest and the induction of a pro-inflammatory senescence-associated secretory phenotype (SASP) that is triggered through an unknown mechanism. Here, we show that during oncogene-induced senescence (OIS), the Toll-like receptor TLR2 and its partner TLR10 are key mediators of senescence *in vitro* and in murine models. TLR2 promotes cell cycle arrest by regulating the tumour suppressors p53-p21^CIP1^, p16^INK4a^ and p15^INK4b^, and regulates the SASP through the induction of the acute-phase serum amyloids A1 and A2 (A-SAA) that, in turn, function as the damage associated molecular patterns (DAMPs) signalling through TLR2 in OIS. Finally, we found evidence that the cGAS-STING cytosolic DNA sensing pathway primes TLR2 and A-SAA expression in OIS. In summary, we report that innate immune sensing of senescence-associated DAMPs by TLR2 controls the SASP and reinforces the cell cycle arrest program in OIS.

## INTRODUCTION

Cellular senescence is a cell cycle arrest program induced by various stresses that renders cells insensitive to mitogenic signals and impairs the proliferation and expansion of damaged cells (*1*). Activation of oncogenes such as *RAS* in somatic cells induces a senescence program termed oncogene-induced senescence (OIS) (*2*). OIS is a cell-intrinsic tumour suppressor mechanism that impairs tumour progression (*2-7*). Senescence is also characterised by the activation of the Senescence-Associated Secretory Phenotype (SASP). The SASP is a cocktail of proinflammatory cytokines, chemokines, growth factors and matrix-remodelling proteins with diverse functions and roles (*8-10*). Through the SASP, senescent cells elicit multiple paracrine effects to promote the normal processes associated with inflammation, wound healing, tissue remodelling and cell plasticity (*11-14*) or it can disrupt tissue homeostasis, promoting ageing and other pathophysiological conditions such as fibrosis and cancer (*12*). The SASP reinforces the cell cycle arrest by activating p53 and the cell cycle inhibitors p21^CIP1^ and p15^INK4b^ (*8, 10, 15*). The SASP also promotes paracrine senescence (*15*) and stimulates immune surveillance that results in clearance of the senescent cells (*16-18*), which collectively contribute to the tumour suppressor program elicited by OIS. Therefore, the immune system can regulate senescence and determine senescent cell fate (*19-21*).

The activation of the SASP is primarily regulated by the transcription factors nuclear factor-κB (NF-κB) and CCAAT/Enhancer Binding Protein Beta (C/EBPβ) (*8, 10, 22*). SASP activation responds to a hierarchical model where interleukin-1α (IL-1α) and interleukin-1β (IL-1β) are the apical cytokines signalling through Interleukin-1 Receptor (IL-1R), triggering a cascade of additional amplifying feedback loops by other cytokines and chemokines of the SASP. As a consequence, NF-κB and CEBPβ activation is sustained and the expression of the SASP is maintained. Interfering with these feedback loops, such as the one set up by IL-1, interleukin-6 (IL-6) or interleukin-8 (IL-8), collapses the network impairing the induction of the SASP and the activation of the cell cycle arrest (*8, 10, 15, 23*).

We have previously shown that the inflammasome regulates the activation of the SASP (*15*). The inflammasome is a multiprotein platform that induces the proteolytic activity of the inflammatory cysteine-aspartic protease Caspase-1 (CASP1). It is a key regulatory component of the innate immune response and the first line of defence against pathogens and damage (*24*). Activation of the inflammasome leads to cleavage and activation of the proinflammatory cytokine IL-1β. Inflammasomes are controlled by a family of receptors called pattern recognition receptors (PRRs). PRRs are receptors of the innate immune system that are activated by interaction with pathogen-associated molecular patterns (PAMPs) or with damage-associated molecular patterns (DAMPs) that are generated endogenously in cells under certain conditions of stress and damage (*25*). There are three major PRR families: Toll-like receptors (TLRs), RIG-I-like receptors (RLRs) and NOD-like receptors (NLRs) (*25*). Upon activation, PRRs induce distinct signal transduction pathways that activate an immune transcriptional program mostly regulated by NF-κB and Interferon Regulatory Factors (IRFs).

Although the steady state signalling of the SASP is relatively well understood, the molecular mechanism(s) that initiates the SASP in OIS and how the inflammasome is primed during OIS remains ill-defined. Here we describe the mechanism that underlies the priming of the inflammasome in OIS by PRRs, identifying the senescence-associated DAMP that initiates the SASP and reinforces OIS.

## RESULTS

### The innate immune receptor TLR2 is induced during cellular senescence

To study OIS, we used the well-characterised IMR90 human diploid fibroblast cell line transduced with an ER:H-RAS^G12V^ fusion protein (henceforth referred to as ER:RAS). ER:RAS oncogenic activity is induced after addition of 4-hydroxytamoxifen (4OHT) to the cultures, resulting in a time controlled OIS response as previously described (*15*) (Fig. 1A). Gene Set Enrichment Analysis of the transcriptome of IMR90 ER:RAS senescent cells showed an enrichment for “innate immunity”, “pattern recognition receptors” and “toll-like receptors (TLRs)” terms (Fig. S1A). Therefore, we decided to explore the role of TLRs in OIS. Analysis of the gene expression of all ten human TLR genes showed a marked induction of Toll-like receptor 2 (*TLR2*) expression in OIS (Fig. 1B). The induction of *TLR2* mRNA correlated with an increase in TLR2 protein (Fig. 1C), which corresponded with the cell cycle arrest during OIS (Fig. 1C). The induction of *TLR2* was also observed by conditioned medium from OIS cells (paracrine senescence) (*15*), after retroviral transduction of oncogenic H-Ras^G12V^, and by the activation of senescence with a DNA damage-inducing agent (Etoposide) (Fig. S1, B to D).

**Figure 1.**
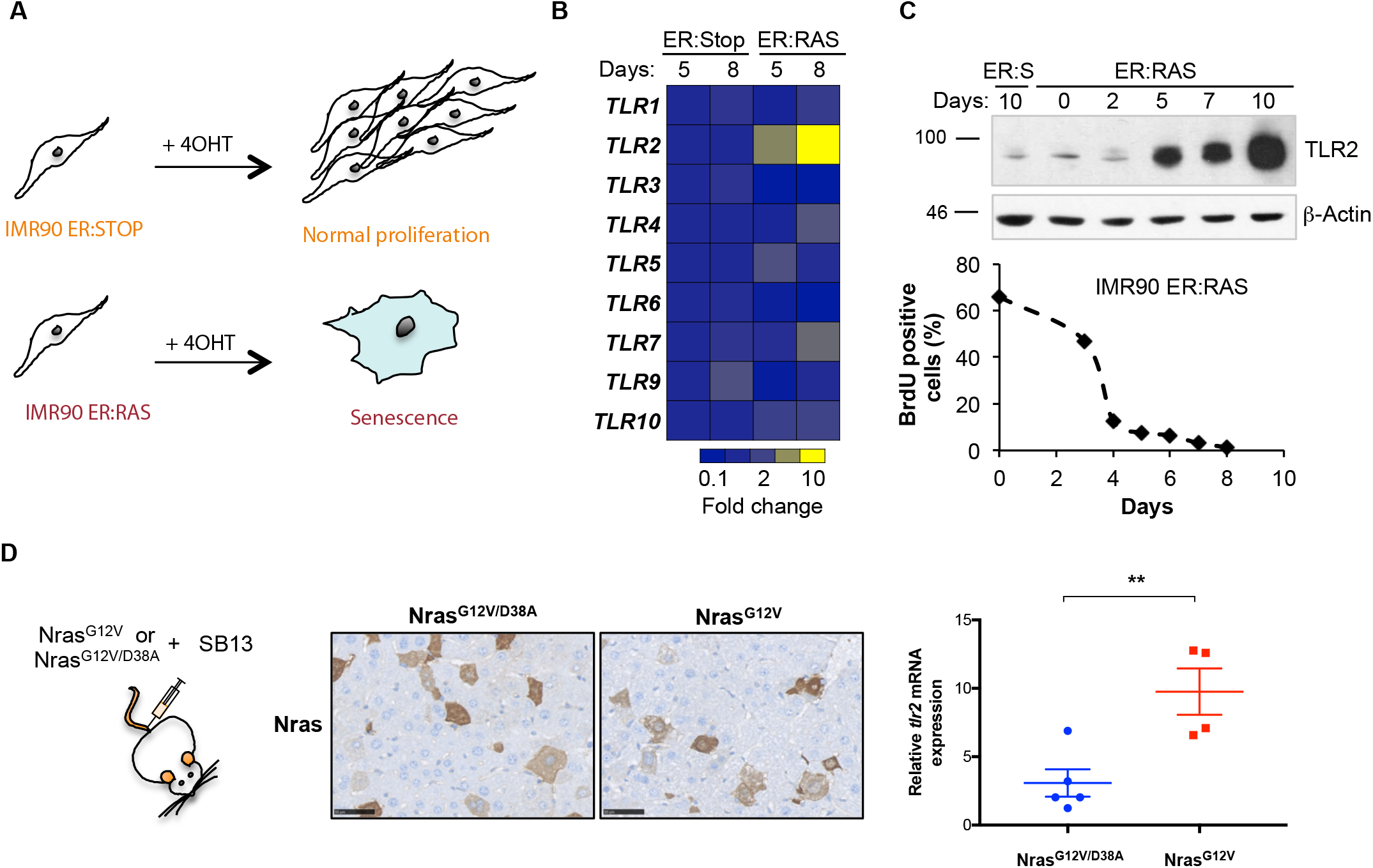
TLR2 expression is induced during Oncogene Induced Senescence. (**A**) Schematic showing IMR90 ER:RAS cells treated 4OHT undergo OIS. IMR90 ER:Stop cells serve as a control and retain proliferative capacity with 4OHT. **(B)** qRT-PCR analysis of TLR family member expression in IMR90 ER:RAS and ER:Stop cells treated with 4OHT for 5 and 8 days. **(C)** Western blot of TLR2 expression in IMR90 ER:RAS and ER:Stop (ER:S) cells with up to 10 days of 4OHT treatment (upper panel). BrdU incorporation in IMR90 ER:RAS cells treated with 4OHT for up to 8 days as indicated. **(D)** Immunohistochemical staining for Nras in liver sections from WT mice 6 days following hydrodynamic delivery of Nras^G12V/D38A^ (n=5) and Nras^G12V^ (n=4) transposons. Scale bar 50µm. RNA was extracted from snap frozen liver samples and *tlr2* mRNA expression was measured using qRT-PCR. Scatter plots represents value per animal with the horizontal line representing group mean ± SEM. Statistical significance was calculated using Students two-tailed *t*-test. **p<0.01.

*In vivo*, we analysed *tlr2* expression in four well-characterized mouse models of senescence. We first examined the expression of *tlr2* in a murine model of OIS in which conditional expression of Kras^G12D^ by Pdx-CRE (KC mouse) induces Pancreatic Intraepithelial Neoplasia (PanIN) (*26*). A marked induction of Tlr2 was observed in ductal pancreatic cells in PanINs with low Ki-67 indexes, indicating that Tlr2 is expressed in early senescent PanINs (Fig. S1E). We then investigated an additional model where OIS is induced in murine hepatocytes via hydrodynamic delivery of a mutant Nras^G12V^ encoding transposon along with a sleeping beauty transposase expressing plasmid (*16*). Plasmid encoding an Nras^G12V^ effector loop mutant, incapable of downstream Nras signalling (Nras^G12V/D38A^), was used as a negative control. Six days after Nras^G12V^ transduction, *tlr2* mRNA expression was significantly increased in comparison to controls (Fig. 1D), which correlated with the expected induction of mRNA expression of the senescence markers *dcr2* and *arf*, and the SASP factor IL-1β (Fig. S1F). We then investigated the expression of *tlr2* in a model of inflammatory-mediated senescence, where knockout of *nfkb1* (*nfkb1^-/-^)* in mice leads to constitutive activation of Nf-κB (*27*). An increased number of cells expressing Tlr2 were detected, which correlated with p21^Cip1^ expression in the lung airways (Fig. S1G). Finally, we observed an increase in the number of cells positively expressing Tlr2 in alveolar cells of the lung in aged mice (Fig. S1H). These results indicate that TLR2 is induced in cellular senescence.

### TLR2 controls the activation of the SASP in OIS

We previously identified that activation of IL-1R signalling by the inflammasome was essential for the induction of the SASP (*15*). However, prior to inflammasome activation, the expression of IL-1β should be primed (*28, 29*). TLR2 forms functional heterodimers with three other Toll-like receptors, TLR1, TLR6 and TLR10 (*30-32*). To evaluate the activity of all four members of the TLR2 network in IL-1β priming, we knocked-down their expression during OIS using four pooled siRNAs (Fig. S2A). While knockdown of TLR1 and TLR6 marginally decreased the expression of IL-1β, TLR2 and TLR10 knockdown strongly decreased its induction (Fig. 2, A to C and fig. S2B,), suggesting a predominant role for these receptors in priming the inflammasome in OIS. To rule out off-target effects from the siRNAs, we tested all four individual siRNAs from the TLR2 and TLR10 pool. Each siRNA produced efficient TLR2 knockdown in IMR90 ER:RAS cells (Fig. S2C) and decreased the expression of IL-1β in OIS (Fig. S2D). Similarly, two single siRNA from the pooled siRNA also efficiently reduced the expression of TLR10 and impaired IL-1β transcriptional activation (Fig. S2, C and E). Importantly, TLR2 and TLR10 knockdown resulted in reduced production of mature active IL-1β (Fig. 2B), and a decrease in the accumulation of mature IL-1β in conditioned media of IMR90 ER:RAS cells (Fig. S2B), indicating a crucial role for the TLR2 network for the inflammasome function in senescence. Inflammasome activation is a key step in the induction of the SASP during OIS (*15*). Therefore, we studied the effect of TLR2 signalling in SASP induction. Targeting TLR2 and TLR10 with siRNA impaired the induction of mRNA expression of the SASP components IL-1α, IL-6, IL-8, CCL20, MMP1, MMP3, INHBA (Fig. 2C) and blocked the induction of IL-8 and IL-6 protein (Fig. 2B and fig. S2F). Moreover, knockdown of TLR2 and TLR10 in IMR90 cells impaired the induction of IL-1β, IL-6, and IL-8 when exposed to conditioned media from senescent IMR90 ER:RAS cells, indicating a contribution of TLR2 in the propagation of the SASP during paracrine senescence (Fig. S2G).

**Figure 2.**
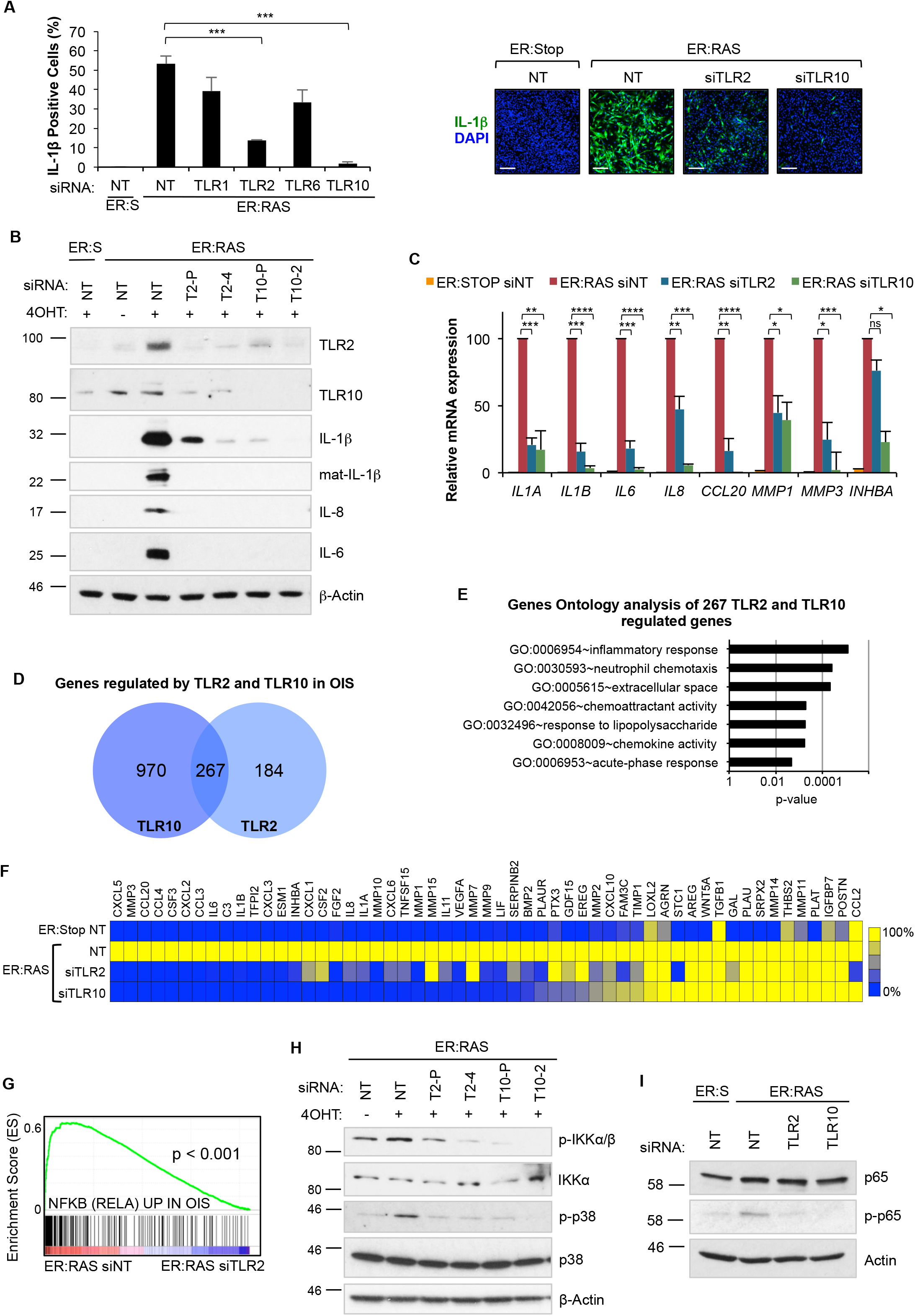
TLR2 and TLR10 regulate the SASP in Oncogene Induced Senescence. **(A)** Immunofluorescence staining and high content analysis for IL-1β expression in IMR90 ER:RAS cells treated with 4OHT for 8 days and repeatedly transfected with pooled siRNA targeting TLR1, TLR2, TLR6 and TLR10. Non-target pooled siRNA (NT) was used as control. Representative images are shown. Scale bar 250 µm. **(B)** Western blot analysis against indicated antibodies in IMR90 ER:RAS cells treated with 4OHT for 8 days and repeatedly transfected with pooled and individual siRNA targeting TLR2 and TLR10. T2-P, siRNA TLR2 pool; T2-4, individual TLR2 siRNA; T10-P, siRNA TLR10 pool; T10-2, individual TLR10 siRNA. Non-target pooled siRNA (NT) was used as control. Western blot against b-Actin is shown as a loading control. **(C)** SASP factor regulation by qRT-PCR in ER:RAS cells treated with 4OHT for 8 days and repeatedly transfected with pooled siRNA targeting TLR2 and TLR10. Results expressed as mean ± SEM of 3 independent experiments. **(D)** Venn diagram shows the number of genes that are significantly induced by TLR2 and TLR10 during OIS, and the intersection represents the number of genes regulated by both TLR2 and TLR10. This signature of 267 genes will be used for gene set enrichment analysis in additional senescence transcriptomes in Fig. 4 and Supplemental Fig. S4 and S7). **(E)** Top regulated terms identified through of coregulated genes in (H) using DAVID gen ontology analysis. Chart bars represents Benjamin adjusted p-value of term enrichment. **(F)** Heat-map of SASP factor expression obtained from the transcriptome analysis (Ampliseq) in IMR90 ER:RAS cells following siRNA knockdown of TLR2 and TLR10 for 8 days of 4OHT treatment. **(G)** GSEA enrichment plot of RELA signature in TLR2 siRNA transfected IMR90 ER:RAS 4OHT induced cells. **(H)** IMR90 ER:RAS cells were transfected with indicated siRNA for 8 days with 4OHT. Western blot for phosphorylation and total levels of IKK α/β and p38 MAPK was performed. **(I)** IMR90 ER:RAS cells were treated with 4OHT and repeatedly transfected with indicated pooled siRNA and non-target (NT) siRNA as control for 5 days. Western blots were conducted for phosphorylation of p65 and total p65 protein levels.

We then performed a transcriptomic analysis of over 20,000 genes of IMR90 ER:RAS cells transfected with siRNA targeting TLR2 and TLR10 (Fig. S2H). We found that TLR2 knockdown significantly regulated more than 1,000 genes, while TLR10 knockdown regulated up to 2,500 genes (Fig. S2H). From the total number of genes positively regulated by TLR2 and TLR10 in OIS, 267 were common targets, indicating a high degree of overlap between genes positively regulated by TLR2 and TLR10 (Fig. 2D). Gene ontology analysis of the genes commonly regulated by TLR2 and TLR10 in OIS showed enrichment for terms related to “inflammatory response”, “chemotaxis", "chemokine activity" and "extracellular space" (Fig. 2E). Moreover, the analysis revealed that TLR2 and TLR10 regulated the expression of most of the proinflammatory SASP, most notably chemokines, cytokines, and metalloproteases (Fig. 2F). TLR2 signals through NF-κB and p38 MAPK (*33*), which are key SASP regulators. A significant enrichment in genes regulated by NF-κB in OIS (*22*) was observed in the transcriptome of control cells compared to TLR2 depleted cells (Fig. 2G), indicating an essential role for TLR2 signalling in NF-κB activation in senescence. To confirm this, we targeted TLR2 and TLR10 with siRNA in 4OHT treated IMR90 ER:RAS cells, and showed a decrease in NF-κB pathway activation (reduced IKKα/β and p65 phosphorylation) (Fig. 2H and I). We also observed a marked decrease in p38 MAPK phosphorylation in TLR2 and TLR10 depleted cells in OIS (Fig. 2H). These results show that TLR2 signalling is a significant contributor to the activation of the SASP by controlling NF-κB and p38 MAPK signal transduction pathways. Altogether, these data indicate that TLR2 signalling is necessary for the induction of the SASP.

### TLR2 reinforces the cell-cycle arrest program in OIS

Activation of the SASP reinforces the cell cycle arrest program in senescence (*8, 10*). Moreover, p38 MAPK is a significant regulator of OIS, controlling the activation of p53 and p16^INK4a^ tumour suppressor genes (*34*). Inhibition of p38 MAPK but not NF-κB with chemical compounds bypassed the proliferative arrest and impaired p21^CIP1^ activation in IMR90 ER:RAS cells, showing a role for p38 MAPK in the reinforcement of the cell cycle arrest in OIS (Fig. S3, A and B). Given the integral role TLR2 plays in the regulation of the SASP and p38 MAPK, we explored whether TLR2 also controls the cell cycle arrest in OIS. Indeed, overexpression of TLR2 induced cell cycle arrest and an increase in the number of SA-β-Gal positive cells (Fig. 3, A and B). Analysis of proliferation after ER:RAS activation showed that targeting TLR2 and TLR10 with siRNA strongly reduced the cell cycle arrest associated with OIS (Fig. 3C), and increased the long term growth of ER:RAS cells (Fig. 3D and fig. S3, C and D). This effect correlated with a decrease in the number of cells positive for SA-β-Gal activity 10 days after ER:RAS activation (Fig. 3E). Importantly, the control of proliferation by TLRs was specific for TLR2 and TLR10, as knockdown of the other eight members of the human TLR family did not show bypass of the cell cycle arrest program (Fig. S3E). Suppression of TLR2 and TLR10 resulted in a reduction of p53 protein levels (Fig. 3G) and a decrease in p21^CIP1^, p16^INK4a^ and p15^INK4b^ mRNA expression (Fig. 6F). Also, TLR10 knockdown induced escape of cell cycle arrest in cells subjected to conditioned media from OIS cells, suggesting a role in cell cycle arrest during paracrine senescence (Fig. S3F). Altogether, these results indicate that TLR2 contributes to the cell cycle arrest program induced during OIS.

**Figure 3.**
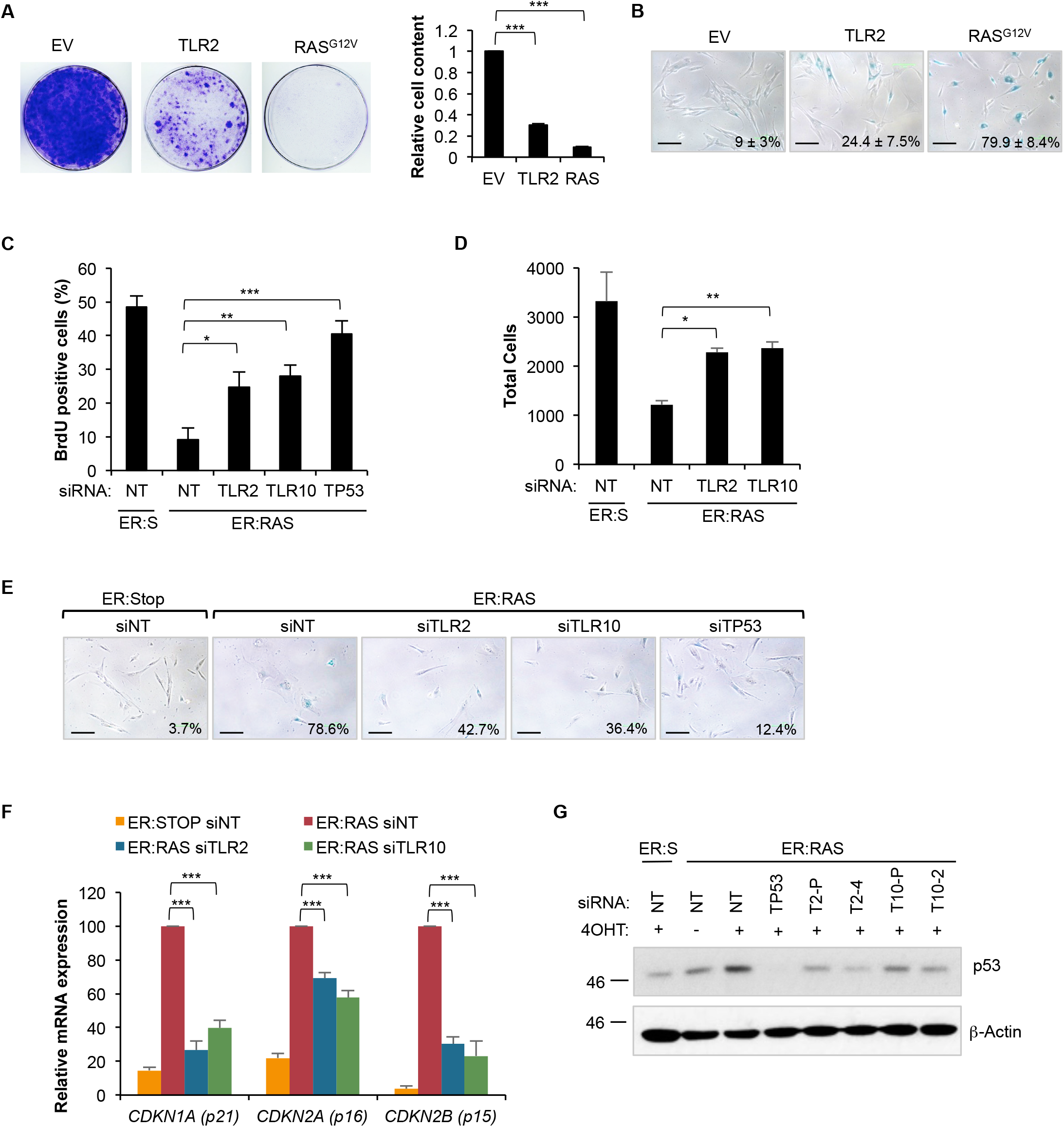
TLR2 reinforces the cell cycle arrest in OIS. **(A)** IMR90 cells infected with TLR2 or H-Ras^G12V^ expression vectors, or empty vector control (EV) were seeded at low density and stained with crystal violet after 2 weeks. The staining was quantified to obtain relative cell content. Results expressed as mean ± SEM of 3 independent experiments. **(B)** SA-β-GAL staining was carried out on TLR2 and HRas^G12V^ expressing cells. Results expressed as mean (% positive cells) ± SEM of 3 independent experiments. **(C-G)** IMR90 ER:RAS cells were treated with 4OHT and repeatedly transfected with indicated siRNA and pooled non-target (NT) siRNA control. siTP53 was used as a positive control. **(C)** After 5 days of treatment, a BrdU incorporation assay was conducted. Results expressed as mean ± SEM of 3 independent experiments. **(D)** Total DAPI stained nuclei counted by high content analysis at 8 days. Results expressed as mean ± SEM of 3 independent experiments. **(E)** After 10 days SA-β-GAL activity assay was conducted. Scale bar 100 mm. **(F)** qRT-PCR analysis of *CDKN1A, CDKN2A* and *CDKN2B* transcripts. Results expressed as mean ± SEM of 3 independent experiments. **(G)** Western blot for p53 expression at eight days. All statistical significance was calculated using One-Way ANOVA. ***p < 0.001, **p < 0.01, * p < 0.05, ns, non-significant.

### TLR2 controls the expression of acute-phase serum amyloid A1 and A2, which are components of the SASP

Gene Set Enrichment Analysis of the transcriptome of TLR2- and TLR10-targeted cells revealed “acute phase response” and “positive regulation of acute inflammatory response” as the top gene ontology terms (highest enrichment score and significance) for both conditions (Fig. S4A). The acute phase response is a clinical indication of an inflammatory event such as infection, trauma or neoplasia, and is characterised by the production and secretion of acute phase proteins, including C-reactive protein and acute phase serum amyloids by the liver (*35*). The two genes most strongly down-regulated by TLR2 and TLR10 knockdown were acute-phase Serum Amyloid A 1 and 2 (*SAA1* and *SAA2*, A-SAA from now on) (Fig. 4A). Other genes annotated as “acute phase response” such as IL-1α, IL-1β, IL-6, SERPINA3, ASS1 and PTGS2 also showed strong regulation by TLR2 and TLR10 in OIS (Fig. 4A). Further mRNA analysis by qRT-PCR of SAA1, SAA2, SERPINA3, PTGS2, STAT3 and IL-6R expression confirmed these results (Fig. 4B), indicating that the expression of A-SAAs is highly induced during OIS and is dependent on TLR2 and TLR10. A time course analysis of OIS revealed that A-SAA expression is induced 5 days after the activation of oncogenic RAS with 4OHT, which overlaps with TLR2 induction (Fig. 3C, 1C and fig. S4B). Treatment of IMR90 cells with Pam2CSK4 (a synthetic agonist for TLR2) induced A-SAA expression, which was increased by one order of magnitude by ectopic overexpression of TLR2 (Fig. S4C), suggesting a role for TLR2 sensing in the induction of A-SAA in OIS. Western blot of the conditioned medium from IMR90 ER:RAS cells showed an accumulation of A-SAA, which was reduced following siRNA knockdown of SAA1 and SAA2 (Fig. 4D fig. S4D), suggesting that A-SAAs are components of the SASP. We then studied the induction of ASAAs in senescence *in vivo*. mRNA expression of saa2 was induced in the lung of *nfkb1* knockout mice in parallel with the senescence markers cdkn1a (p21) and cdkn2a (p16) (Fig. S4E). Moreover, increased expression of Saa1 was detected in early, low proliferative PanINs in KC mouse (Fig. S4F). In summary, these results show that A-SAAs are components of the SASP regulated by TLR2.

**Figure 4.**
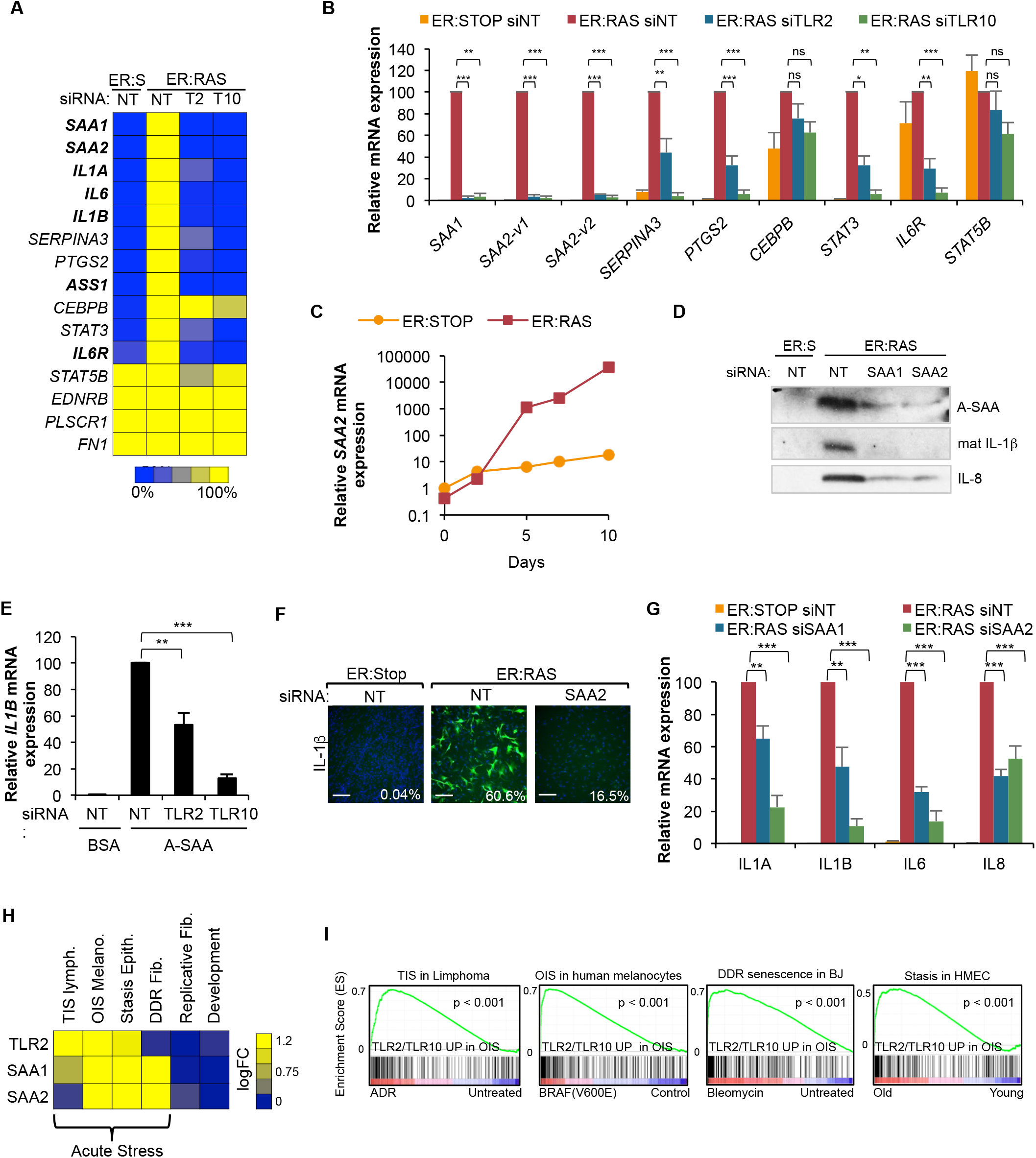
TLR2 and TLR10 regulate the activation of genes of the acute phase response during OIS. **(A)** Heat-map showing the relative fold change of acute phase response transcripts of samples from the “acute phase response” gene set from the mRNA transcriptomes. Transcriptome analysis (Ampliseq) was performed in mRNA from IMR90 ER:RAS cells transfected with pooled siRNA for TLR2 and TLR10, and non-target pool (NT) as a control. Genes with significant changes between NT siRNA control and both TLR2 and TLR10 knockdown are in bold characters. Adjusted p-values were calculated using Benjamini and Hochberg (BH) false discovery rate of 3 independent experiments. Bold genes represent adjusted p-value < 0.05. **(B)** qRT-PCR validation of acute phase response targets from samples obtained similarly to (A). Results expressed as mean ± SEM of 3 independent experiments. Statistical significance was calculated using One-Way ANOVA and Dunnett’s multiple comparison’s tests. ***p < 0.001, **p < 0.01, * p < 0.05, NS, non-significant. **(C)** qRT-PCR analysis of A-SAA expression in IMR90 ER:RAS and ER:Stop cells with up to 10 days of 4OHT treatment. **(D)** IMR90 ER:RAS cells were treated with 4OHT and repeatedly transfected with pooled siRNA targeting SAA1 and SAA2, and non-target (NT) siRNA as control for 8 days. Western blot of the conditioned medium for indicated antibodies. **(E)** IMR90 cells transfected with pooled siRNA for TLR2 and TLR10 were treated with 10 µg/ml A-SAA for 3 hours and qRT-PCR was performed to measure *IL1B* expression. Results are expressed as mean ± SEM of 3 independent experiments. **(F)** Immunofluorescence staining and quantification of IL-1β expression by high content analysis. Scale bar 250 µm. **(G)** qRT-PCR for *IL1A*, *IL1B, IL6, IL8* expression. Results expressed as mean ± SEM of 3 independent experiments. **(H)** Heatmap showing TLR2 SAA1 and SAA2 expression in available transcriptomic data from Therapy-induced senescence lymphoma cells (TIS) (GSE31099), Oncogene-induced senescence mediated by mutant BRAF in human melanocytes (OIS) (GSE46801), Stasis in HMEC (GSE16058), DNA damage induced senescence in BJ cells (DDR) (GSE13330), Replicative senescence in BJ cells (replicative) (GSE13330) and developmental senescence in the mesenephos (developmental) (GSE49108). **(I)** Gene-set enrichment analysis plots for the 267 genes regulated co-regulated by TLR2 and TLR10 in OIS (Supplemental Fig. S2H) in the transcriptomes from (I). All statistical significance was calculated using One-Way ANOVA. ***p < 0.001, **p < 0.01, * p < 0.05, ns, non-significant.

### A-SAAs are potent senescence-associated DAMPs that control the SASP through TLR2 signaling

Previous reports have shown that A-SAAs have cytokine-like activity by interacting with TLR2 (*36*). Thus, we decided to explore the role of A-SAAs regulating the SASP in OIS. Recombinant A-SAA (rA-SAA) induced IL-1β mRNA expression with a similar effect to that achieved with an equimolar dose of the synthetic TLR2 agonist Pam2CSK4 (Fig. S5A). Also, TLR2 overexpression in IMR90 cells enhanced rA-SAA-dependent induction of the SASP (Fig. S5, B and C). Interference of TLR2 function with neutralising antibodies (Fig. S5D) or with siRNA against TLR2 and TLR10 (Fig. 4E, fig. S5E) also inhibited rASAA-mediated induction of SASP components. These results suggest that A-SAAs directly activate TLR2, resulting in priming of the inflammasome and SASP induction. To further confirm this, we decided to target the expression of A-SAAs during OIS with siRNA, showing that the induction of IL-1β and the SASP was impaired (Fig. 4, F and G). Moreover, inactivation of A-SAA during OIS impaired the accumulation of mature IL-1β and IL-8 in conditioned media (Fig. 4D). Altogether, these data indicate that A-SAAs are the DAMPs that signal through TLR2 to prime the inflammasome and regulate the SASP in OIS.

To assess the relative importance of TLR2 and A-SAA mediated SASP regulation, we compared the effect of TLR2, TLR10 and SAA2 with other previously described regulators of the SASP, including: the DDR pathway (ATM), the cGAS-STING cytosolic DNA sensing pathway, the master SASP-regulator IL-1 (IL-1R), the mTOR pathway, and the essential SASP transcription factors NF-κB (RELA) and CEBPβ (*37*). Knockdown of the A-SAA-TLR2 pathway had a similarly negative effect on IL-1β induction to that following knockdown of the other known SASP regulators (Fig S5, F and G)). Also, knocking-down the other A-SAA receptor TLR4 did not decrease IL-1β induction, suggesting a specific role for A-SAA-TLR2 signalling in OIS (Fig. S5G).

Finally, we wanted to assess the existence of this regulatory pathway in additional cell types and senescence triggers. We first explored the expression of TLR2, SAA1 and SAA2 in available transcriptomic datasets of senescence (Fig. 4H). This analysis revealed that the A-SAA-TLR2 pathway was induced in senescence activated by acute stresses such as Therapy Induced Senescence (TIS) in lymphoma cells (*38*), BRAF^V600E^ induced senescence in human melanocytes (*39*), stasis in primary human mammary epithelial cells (*40*), and DDR induced senescence in human dermal fibroblasts (BJ) (*41*), while it was not induced in replicative senescence (*41*) and programmed developmental senescence in the murine mesonephros (*42*)(Fig. 4H). Strikingly, the gene set composed by the 267 genes co-regulated by TLR2 and TLR10 in OIS (Fig. 2D) was significantly enriched (p<0.01) only in those transcriptomes with activation of A-SAA-TLR2 expression (Fig. 4I), while this correlation was not found with the transcriptome of replicative and developmental senescence (Fig. S5H), suggesting a role for the A-SAA and TLR2 pair in senescence induced by several acute stresses in distinct cell types. Altogether, these results indicate that A-SAAs and TLR2 establish a master innate immune pathway with a key role in the control of the transcriptome in senescence.

### The cGAS-STING cytosolic DNA sensing pathway controls the induction of TLR2 and A-SAA in OIS

Experiments targeting RELA and ATM revealed that the activation of TLR2 and ASAA is dependent on NF-κB and the DDR in OIS (Fig 5A). Recently, it has been shown that in senescence, NF-κB is activated by the cGAS-STING cytosolic DNA sensing pathway in response to the accumulation of cytoplasmic chromatin fragments released from the damaged nucleus (*43*). Thus, we speculated that this pathway could be responsible for the induction of TLR2 and A-SAA in OIS. Inactivation of cGAS and STING with siRNA in OIS strongly impaired the transcriptional activation of TLR2, SAA1 and SAA2 mRNA expression (Fig. 5B), indicating a role for the DNA sensing pathway in the regulation of TLR2 and A-SAA. Moreover, direct activation of cGAS-STING by transfecting dsDNA into IMR90 cells induced the formation of STING homo-dimers, which is a hallmark of its activation, and induced TLR2 expression, which was reduced by STING knockdown (Fig. 5, C and D). Finally, knockdown of TLR2 and TLR10 did not reduce the formation of STING homo-dimers in OIS (Fig. 5D), suggesting that TLR2 signalling is downstream of the cGAS-STING pathway. Altogether, these results indicate that A-SAA and TLR2 are downstream of the cGAS-STING cytosolic DNA sensing pathway in OIS.

**Figure 5.**
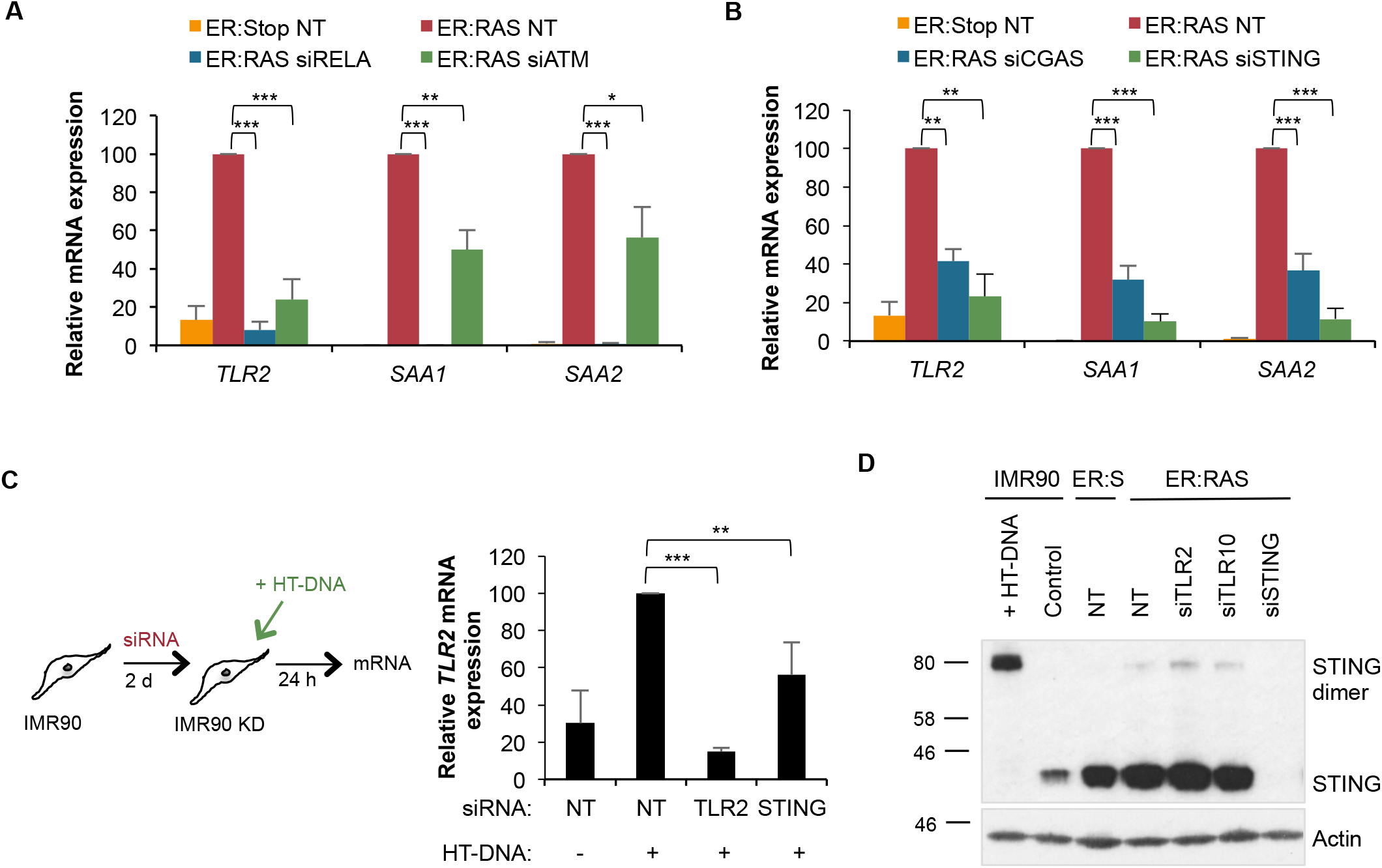
A-SAA and TLR2 expression is dependent upon STING activation. (**A, B**) IMR90 ER:RAS cells were treated with 4OHT and repeatedly transfected with indicated pooled siRNA and non-target (NT) siRNA as control for 8 days. *TLR2, SAA1* and *SAA2* transcripts were measured by qRT-PCR. Results expressed as mean ± SEM of 3 independent experiments. **(C)** IMR90 cells were transfected with siRNA targeting TLR2 and STING for 2 days followed by transfection with 2.5 µg herrings-testes DNA (HT-DNA) for 24 hours. *TLR2* transcripts were measured by qRT-PCR. Results are expressed as mean ± SEM of 3 independent experiments. **(D)** Western blot for STING dimerization. HTDNA transfection of IMR90 cells were used as positive control for STING dimerization. IMR90 ER:RAS cells were transfected with siRNA targeting TLR2, TLR10 and STING for 8 days with 4OHT. All statistical significance was calculated using a One-Way ANOVA. ***p < 0.001, **p < 0.01, * p < 0.05, ns, non-significant.

### TLR2 regulates the SASP *in vivo*

The strong enrichment of the TLR2/TLR10 OIS dependent gene set in the publically available transcriptome of Kras^G12D^ driven PanINs in KC mice (*44*), suggested a role for TLR2 in this *in vivo* model (Fig. S6A). Therefore, we decided to test in KC mice the effect of knocking-out *tlr2* (*tlr2^-/-^)* on SASP activation. We observed a reduction in IL-1α staining in PanIN epithelial cells of *tlr2^-/-^* mice compared to wild-type (WT) mice (Fig. 6A), indicating a causal role for TLR2 in the activation of the SASP in PanIN. We also assessed the effect on SASP expression after hydrodynamic delivery of Nras^G12V^ transposons into hepatocytes of *tlr2^-/-^* mice (Fig. 6, B and C). As expected, the transduction of oncogenic Nras^G12V^ induced the accumulation of IL-1β positive cells (Fig. 6B) and the induction of IL-1β, IL-1α and IL-6 mRNA expression (Fig 6C) in the liver of WT mice when compared to the inactive mutant (Nras^G12V/D38A^) control. In contrast, Nras^G12V^ failed to induce IL-1β positive cells or mRNA expression in *tlr2^-/-^* mice (Fig. 6, B and C). Moreover, the induction of the other SASP components IL-1α and IL-6 was also impaired in *tlr2^-/-^* mice (Fig. 6C). In summary, these results indicate that TLR2 is a necessary for SASP induction during OIS *in vivo*.

**Figure 6.**
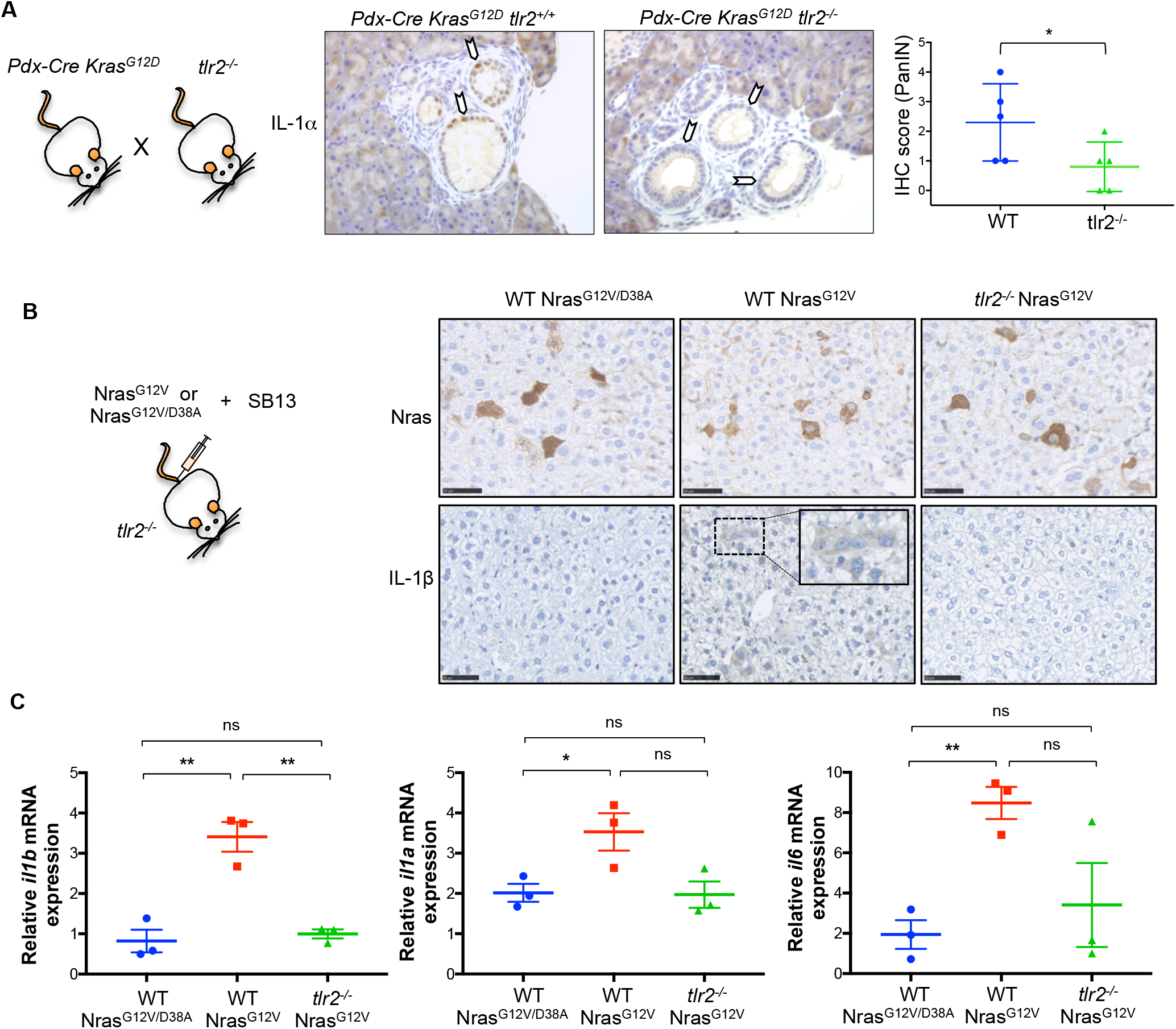
*tlr2* is necessary for SASP activation *in vivo*. **(A)** Representative immunohistochemical (IHC) staining and IHC score quantification of IL-1α in PanIN generated in *tlr2^+/+^* or *tlr2^-/-^ Pdx-Cre Kras^G12D^* mice. Scatter plot represents value for individual animals (dots) and the horizontal line represents group mean (n=5) ± SEM. Statistical significance was calculated using one-tailed students *t*-test. *p<0.05 **(B)** Representative IHC staining for Nras (top row) and IL-1β (bottom row) in liver sections from wild-type (WT) and *tlr2^-/-^* mice 6 days after receiving hydrodynamic delivery of Nras^G12V/D38A^ negative control or oncogenic Nras^G12V^ transposon as indicated. Image magnification is shown in the il-1β box insert. Scale bar 50µm **(C)** qRT-PCR results for SASP factors IL-1β, IL-1α and IL-6 from liver samples from corresponding mice in (B). Scatter plot represents value for individual animals (dots) and the horizontal line represents group mean (n=3) ± SEM. Statistical significance was calculated using two-tailed students *t*-test. *p<0.05, **p<0.01. ns, non-significant.

## DISCUSSION

We describe here a new and essential innate immune signalling pathway in OIS established between TLR2 and A-SAAs that initiate the SASP and reinforce the cell cycle arrest (Fig. S6B). Importantly, we identify new important SASP components, A-SAAs, which are the senescence-associated DAMPs sensed by TLR2 after oncogenic stress *in vitro* and *in vivo*. Therefore, we are reporting that innate immune sensing is critical in senescence.

We propose that cellular senescence shares mechanistic features with the activation of innate immune cells, and could be considered a program of the innate immune response by which somatic cells switch their regular role to acquire an immune function under certain conditions of stress and danger, for instance upon oncogene activation. In OIS, super-enhancer elements regulated by BRD4 adjacent to the SASP have been shown to regulate the immune surveillance of senescent cells (*45*). In the same study, it was demonstrated that TLR pathways in general, and TLR2 in particular, were associated with activation of typical enhancers during OIS, suggesting that enhancer remodelling might shape the switch that activates this immune sensing program during senescence. Additionally, cytosolic DNA sensing by the innate immune sensor cGAS has also been identified as an essential step in the activation of the SASP (*43, 46*). In our study, we have shown that the induction of TLR2 and A-SAA depends on the activation of cGAS and STING in senescence, and they appear to function downstream of STING to regulate the SASP and NF-κB (Fig. S6B). Future research will be necessary to understand further the crosstalk between both pathways in senescence. For example, Dou et al. proposed that p38 MAPK signalling impairs the activation of the interferon response arm of the cGAS-STING pathway, prioritising the NF-κB response. Importantly, we observed that TLR2 is essential for both the steady-state activation of p38 MAPK and NF-κB during OIS. Therefore, it is tempting to speculate that TLR2 may interact with the cGAS-STING pathway in senescence by actively impairing the activation of interferon response genes. As a result of these net interactions between innate immune receptors, TLR2 signalling is critical for the SASP and the cell cycle arrest, and coincides with the role of both p38 MAPK and NF-κB signal transduction pathways in OIS (*8, 10, 22, 34*).

Importantly, besides revealing a role for TLR2 in SASP induction and cell cycle regulation, we identified the DAMP that activates TLR2 in OIS. Acute phase proteins SAA1 and SAA2 act to prime the TLR2-mediated inflammasome, and in turn, their full induction depends on TLR2 function. Hence, they establish a foundational feedback loop that controls the SASP (Fig. S6B). A-SAAs are systemically produced in the liver and released into the bloodstream during an acute inflammatory response (*35*). Our identification of these molecules as mediators of senescence suggests that systemic elevation of A-SAAs might have an impact on the accumulation of senescent cells and the activation of their proinflammatory program at the organismal level. We found activation of TLR2 expression in parallel to A-SAA in models of OIS in mice, in inflammation-induced senescence, in ageing and in different *in vitro* systems of senescence. Also, we have shown that TLR2 controls the activation of the SASP *in vivo*. Moreover, we have observed a dose-dependent effect for TLR2 in A-SAA sensing and a role for TLR2 in SASP activation during paracrine senescence. Altogether, these data suggest that systemic A-SAA elevation during acute inflammation could impact upon cells expressing TLR2, thereby promoting ageing and other pathological roles of senescence. Further investigation may reveal additional physiological circumstances under which senescence is induced or reinforced by the interaction of TLR2 with A-SAA, or indeed with other endogenous DAMPs or exogenous PAMPs from the microbiome. Such circumstances could have implications for organismal wellbeing, in particular, the development of ageing and cancer.

Finally, in recent years several strategies have been implemented to eliminate senescent cells or to modulate the activation of the SASP in anti-ageing and cancer therapies (senotherapies) (*47-49*). For example, genetic targeting for the elimination of senescent cells can delay organismal ageing and ageing-associated disorders (*50, 51*). Furthermore, the pharmacological suppression of the SASP has been shown to improve homoeostasis in tissue damage and ageing (*49*). However, most of these manipulations are directed to essential homeostatic regulators such as mTOR or crucial proinflammatory mediators such as IL-1 signalling. Here we propose the alternative of manipulating A-SAATLR2 as a new rationale for anti-SASP therapies aiming to manipulate non-essential and senescence specific signalling pathways.

## MATERIAL AND METHODS

### Cell Culture

HEK293T and IMR90 female human foetal lung fibroblast cells were obtained from ATCC. All cells were maintained in Dulbecco’s Modified Eagle’s Medium (Thermo Fisher Scientific) supplemented with 10% Foetal Bovine Serum (Thermo Fisher Scientific) and 1% antibiotic-antimycotic solution (Thermo Fisher Scientific). IMR90 cells were kept at 5% CO_2_, ambient O_2_ at 37°C. All cell lines were regularly tested for mycoplasma contamination using the Mycoalert Mycoplasma Detection Kit (Lonza). Cell counting and viability were performed using the Muse^®^ Count & Viability Assay Kit in a Muse Cell Analyser (Merck Millipore).

### Experiments with mice

All work was compiled with the UK Home Office guiding principles for the care and use of laboratory animals. Mice carrying a conditional *Pdx1–Cre Kras^G12D/+^* allele were used and have been described previously (*26*). Aging experiments were carried out on male wild-type C57BL/6 mice or male *nfkb1^-/-^* mice on a pure C57BL/6 background at 6.5, 9.5 and 24 months of age. For hydrodynamic tail vein injection experiments *tlr2^-/-^* mice on a C57BL/6 background were purchased from the Jackson Laboratory (JAX^®^). Male and female *tlr2^-/-^* mice and wild-type siblings, aged between 8-12 weeks were included in the study. Plasmids for hydrodynamic injection were prepared using the QIAGEN plasmid maxi kit (QIAGEN, Germany) as per the manufacturers instructions. Animals received 6 µg of a sleeping beauty transposase encoding plasmid (CMV-SB 13 transposase) and 20 µg of NRas^G12V^/GFP encoding plasmid (pT3-NRas^G12V^-IRES-GFP) diluted in physiological saline to 10% of the animals body weight (approximately 2ml), delivered via the lateral tail vein within 10 seconds. Plasmid encoding an NRas^G12V^ effector loop mutant (pT3-NRas^G12V/D38A^-IRES-GFP), incapable of downstream NRas signalling, was used as acontrol. Mice were culled after 6 days and liver tissue was harvested.

### Chemical compounds and neutralising antibodies

Oncogene-induced senescence was induced by treating IMR90 ER:RAS cells with 100 nM 4-hydroxytamoxifen (4OHT) (Sigma). IMR90 ER:STOP are used as a negative control. For DNA damage-induced senescence, IMR90 cells were treated with 100 µM Etoposide for 48 hours. TLR2 was blocked by incubating cells with 10 µg/ml of anti-TLR2 (R&D, MAB2616) or IgG2B Isotype control. Chemical inhibitors used were 10 µM BAY-117082, 10 µM SB202190, 10 µM JAK inhibitor I, 10 µM STAT3 inhibitor V and 10 µM IRAK1/4 inhibitor (all Calbiochem). TLR2 in IMR90 cells was primed with 10 µg/ml Recombinant Human Apo-SAA (Peprotech, #300-30) or 1 µg/ml Pam2SK4 (Tocris, #4637).

### Conditioned Medium for paracrine senescence transmission

For the production of conditional medium used in paracrine senescence, IMR90 ER:STOP or ER:RAS cells were cultured with 100 nM 4OHT in DMEM supplemented with 10% FBS for 4 days, followed by DMEM supplemented with 1% FBS and 4OHT for a further 4 days. The resulting conditioned medium was filtered with 0.2 µm syringe filters (Millipore) and reconstituted with a solution of DMEM supplemented with 40% FBS at a ratio of 3:1.

### Construction of plasmids

TLR2 cDNA was amplified from the pcDNA3-TLR2-YFP plasmid (Addgene 13016) using primers flanked with XhoI sites. Amplified genes were cloned into the MSCV-puro vector. pLN-ER:RAS, LSXN-ER:Stop, MSCV-Ras^G12V,^CMV-VSVG and pNGVL-Gag-Pol vectors were described elsewhere (*15*)

### Retroviral production and infection

For retroviral production, retroviral vectors were co-transfected with VSV-G envelope plasmid and Gag-Pol helper vector using Polyethylenimine Linear, MW 25.000 (PEI) (Polysciences) into HEK293T cells. Viral supernatant was collected from the HEK293T cells two days after transfection and passed through a 0.45 µm (Merck Millipore) syringe filter to eliminate cells. The viral supernatant was supplemented with 4 µg/ml Hexadimethrine bromide (Polybrene) (Sigma) and used to incubate IMR90 cells, with subsequent viral supernatant collection and treatment of IMR90 cells every three hours. After three rounds of infection, the medium was changed to fresh DMEM with 10% FBS and cells were allowed to grow for 2-3 days prior to selection with 1 µg/ml puromycin (Invivogen) for a further 7 days before seeding for indicated experiments.

### Immunohistochemistry

Formalin fixed, paraffin embedded sections were dewaxed, rehydrated through graded ethanol solutions (100, 90, and 70%) and washed in distilled H_2_O. Endogenous peroxidase activity was blocked by immersing sections H_2_O_2_ (Sigma, H1009). To retrieve antigens, sections were boiled in 0.01 M citrate (pH 6.0). Sections were incubated with the primary antibody overnight at 4°C (Antibody information is provided in the supplementary table). For the Pdx1-Cre *KrasG12D/+* mice experiment, EnVision^TM^ +Dual Link system-HRP (DAB+) kit (K4065, Dako), was used for Ki67 staining. The total number of Ki67 positive cells per PanIN, and the total cells per PanIN were counted, and thus the percentage of Ki67 positive cells per PanIN was calculated. The mean score for each mouse was calculated and these scores were plotted on a box plot. Consecutive sections were stained with antibodies against Tlr2 and Saa1. For the *nfkb^-/-^* mice experiment, biotinylated secondary antibody was added and detected using the rabbit peroxidase ABC kit (Vector Laboratories, PK-4001), according to the manufacturer’s instructions. Substrate was developed using the NovaRed kit (Vector Laboratories, SK-4800). Staining was analysed with a NIKON ECLIPSE-E800 microscope, and images were captured with a Leica DFC420 camera using the LAS software (Leica). 10-15 random images were captured per section and the percentage of positively stained cells determined from total number of cells before an average per mouse was calculated. For the hydrodynamic tail vein injection experiment, biotinylated secondary antibody were added and detected using the R.T.U Vectastain^®^ kit (Vector, PK-7100). Substrate was developed using the DAB substrate kit (Abcam, ab64238). Staining was analysed with a Hamamatsu NanoZoomer XR microscope, and images were captured using the NDP scan v 3.1 software (Hamamatsu).

### HT-DNA transfection

MR90 cells were seeded into 6 well tissue culture plates, incubated overnight and transfected with 2500ng of herrings-testes DNA (HT-DNA) using Lipofectamine 2000 following the manufacturer’s protocol. Samples collected for analysis 24 hours after transfection.

### siRNA transfection

Reverse siRNA transfection was carried out using 30 nM siRNA (Dharmacon, GE Healthcare). siRNA sequences are provided in supplementary tables. DharmaFECT1 (Dharmacon, GE Healthcare) transfection reagent was diluted in DMEM and added to the siRNA containing wells. This complex was allowed to form whilst cells were trypsinised and prepared for plating. IMR90 ER:RAS or ER:STOP cells (2000 cells/well in 96 well plate; 66,000 cells/well in 6 well plate) were plated into the siRNA containing wells with 100 nM 4OHT for the activation of ER:RAS. Due to the transient nature of siRNA, to maintain the knockdown for up to eight days, the cells were forward transfected with medium containing the same proportions of siRNA, transfection reagent and 4OHT on days 3 and 5.

### Total RNA preparation and quantitative real-time PCR

RNA was extracted from IMR90 cells using the RNeasy Plus kit (QIAGEN) following the manufacturer’s instructions. For mRNA expression in murine livers, snap frozen specimens were homogenized with trizole and RNA was extracted using the QIAGEN RNeasy plus mini kit (QIAGEN, Germany) according to the manufacturers instructions. cDNA was synthesised using qScript cDNA SuperMix (Quanta Biosciences) following the manufacturer’s instructions from 1µg of RNA in a 40µl reaction. RT-qPCR was performed using 1 µl cDNA as a template per reaction well and SYBR^®^ Select Master Mix (Life Technologies) using 200 nM of forward and reverse primers in 20 µl. Samples were run in triplicate on a StepOnePlus Cycler (Thermo Fisher Scientific).

For the nfkb^-/-^ aging experiment, RNA was extracted from solubilised lung tissue using the RNeasy Mini Kit (QIAGEN, 74106) according to the manufacturer’s instructions. cDNA was generated using the Omniscript RT Kit (QIAGEN, 205110) as per the user manual. RT-qPCR was performed using 4 µl cDNA as a template per reaction well using Power SYBR^®^ Green (Invitrogen, 4367659) PCR Master mix plus 100 nM of forward and reverse primers to form a final reaction volume of 10 µl. Samples were run in triplicate in a C1000TM Thermal Cycler, CFX96TM Real-Time System (Bio-Rad) and Bio-Rad CXF manager software.

mRNA expression analysis was carried out using the change in Ct method and normalised to levels of the housekeeping gene actin or ribosomal protein 18S (nfkb^-/-^ experiment only) to obtain relative mRNA expression. All primers used are listed in the supplementary table.

### Immunofluorescence and high content microscopy

All immunofluorescence staining and imaging was performed on the ImageXpress High Content Analysis microscope (Molecular Devices) as previously described (*52*). All steps were carried out at room temperature. Following the indicated treatment, cells were fixed with 4% PFA for 1 hour and permeabilised with 0.2% Triton X-100 for 10 mins. After 3 washes with PBS, the cells were blocked with 0.2% fish-skin gelatin/BSA/PBS for 1 hour. Primary antibodies were diluted in blocking solution as indicated and incubated for 30 mins followed by 3 washes in PBS. Appropriately conjugated Alexa-fluor secondary antibodies were diluted 1:1000 in blocking solution and incubated on the cells for 40 minutes followed by a further three washes in PBS. Finally, 1 µg/ml DAPI was added to the cells for 20 minutes after which the plates underwent a final round of washes before imaging on the ImageXpress microscope. Quantification of immunofluorescence images was conducted using the MetaXpress software (Molecular Devices) as previously described (*52*).

### Western blot analysis

Whole cell lysates were prepared by lysing the cells with an appropriate volume of RIPA buffer (10mM Tris pH 7.4, 100 mM NaCl, 1 mM EDTA, 1 mM EGTA, 0.1% SDS, 1% Triton X-100, 1 mM β-Mercaptoethanol, 0.5% Sodium Deoxycholate, 10% glycerol, phosphatase inhibitor cocktail III, protease inhibitor cocktail V, EDTA free). Lysates were incubated on ice for 15 minutes prior to centrifuging at 14,000 RPM for 20 minutes to clear lysates. Supernatants were collected into a clean tube and quantified using the Bradford assay. Samples were prepared for loading by mixing with Laemmli sample buffer and boiled at 95°C for 5 minutes. Protein samples were resolved by PAGE and transferred onto nitrocellulose membrane using the iBlot Dry Blotting System (Thermo Fisher Scientific). Membranes were blocked for 1 hour in 5% milk/TBS-1% Tween (TBST). Indicated primary antibodies were diluted in 5% milk/TBST or 5% BSA/TBST and incubated at 4°C with gentle agitation overnight. Blots were washed a minimum of 3 times in TBST over 30 minutes. Secondary antibodies were prepared in 5% milk/TBST and membranes incubated at room temperature for 1 hour. Blots were washed as before and then Enhanced Chemical Luminescence (Amersham) detection reagent was applied for detection. Antibody information is provided in the supplementary table.

### Cell proliferation assays

The BrdU incorporation assay was used to measure the number of cells actively replicating DNA. Cells were treated as indicated in 96 well plates and incubated with 10 µM 5-Bromo-2’-deoxyuridine BrdU (Sigma) for 16-18 hours prior to fixation and immunostained as described using anti-BrdU antibody with 0.5 U/μL DNAse (Sigma) in 1mM MgCl containing PBS.

To visualise and quantify long-term growth, cells were plated at low density (50,000 cells/10 cm plate) and maintained for 10-15 days. The cells were fixed using 1% glutaraldehyde (Sigma) for 1 hour and dried overnight prior to staining with 0.15% crystal violet solution for 2 hours. The plates were then washed, dried and scanned for documentation. For quantification, crystal violet was extracted frim stained plates with 1% acetic acid and quantified by absorbance read at 595nm.

### Senescence-associated beta-galactosidase (SA-β-Gal) assay

SA-β-Gal staining solution was prepared using 20X KC (100mM K_3_FE (CN)_6_ and 100 mM K_4_Fe (CN)_6_*3H_2_O in PBS), 20X X-Gal solution (ThermoFisher Scientific) diluted to 1X in PBS/1 mM MgCl_2_ at pH 5.5-6. The cells were treated as indicated and fixed in 0.5% glutaraldehyde (Sigma) for 10 minutes at room temperature. Cells were washed twice in PBS/1 mM MgCl_2_ pH 5.5-6 prior to incubation in 2 ml staining solution for 24 hours at 37°C.

### Ampliseq Transcriptome profiling

RNA samples were assessed for quality on the Agilent Bioanalyser with the RNA Nano chip, providing an RNA Integrity Number (RIN). Samples were quantified using the Qubit^®^ 2.0 fluorometer and the Qubit^®^ RNA Broad Range assay. 10 ng of RNA was reverse-transcribed to make cDNA, and then target genes were amplified for 12 cycles of PCR using the Ion AmpliSeq™ Human Gene Expression Core Panel. This panel contains 20,802 amplicons (41,604 primers) of approximately 150 bases in length in a single pool. Ion Torrent sequencing adapters and barcodes were ligated to the amplicons and adapter-ligated libraries were purified using AMPure XP beads. Libraries were quantified by qPCR and diluted to 100 pM. Templates were prepared using the Ion PI Hi-Q OT2 200 Kit and sequenced using the Ion PI Hi-Q Sequencing 200 Kit. The Ion Proton platform was used to analyse the data. Analysis of gene expression was performed using the software package babelomics 5.0 (*53*) (*54*). Statistical analysis of significance was performed using Benjamini and Hochberg (BH) FDR multiple test. Gene Set Enrichment Analysis (GSEA) was performed using GSEA 3.0 software from the Broad Institute (www.gsea-msigdb.org) (*55*). Gene Ontology Analysis was performed using DAVID functional annotation web resource (https://david-d.ncifcrf.gov).

### Determination of IL-1β in conditioned medium

Supernatant of medium from siRNA treated cells was collected for analysis of IL-1β content. The medium was combined with 6X Laemmli sample buffer and IL-1β expression determined by immunoblotting as described. Conditional medium samples were also analysed for IL-1β expression by ELISA using the Human IL-1β ELISA Ready-Set-Go! Kit (Affymetrix eBiosciences) following the manufacturer’s instructions.

### Quantification and Statistical Analysis

Immunohistochemistry images were quantified using ImageJ analysis software and graphs and statistical analysis carried out using GraphPad Prism version 7.0, GraphPad Software, San Diego California USA.

Further information and requests for resources and reagents should be directed to and will be fulfilled by the Lead Contact, Juan Carlos Acosta.

## AUTHOR CONTRIBUTIONS

Conceptualisation, J.C.A.; Formal Analysis, P.H, F.R.M., N.T., J.B., J.F.P., J.P.M, and J.C.A; Investigation, P.H., F.R.M., I.F.D, N.T., C.J.R, J.B., J.P.M., L. B., and J.C.A.; Resources, J.P.M. L. B., and V.G.B.; Data Curation, J.C.A; Writing-Original Draft J.C.A.; Writing-Review & Editing, A.J.F, F.R.M., and P.H; Visualisation, P.H., I.F.D., N.T., J.B., J.F.P. and J.C.A; Supervision, J.C.A.; Funding Acquisition, J.C.A.

## ACKNOWLEDGEMENTS

This work was supported by Cancer Research UK (C47559/A16243 Training & Career Development Board - Career Development Fellowship), University of Edinburgh (Chancellor’s Fellowship) and the Wellcome Trust-ISSF. P.H., I.F.D and N.T. were funded by the University of Edinburgh Chancellor’s Fellowship. J.F.P and J.B. were funded by BBSRC (grant BB/K017314/1). F.R.M is funded by a Wellcome Trust Clinical Research Fellowship through the Edinburgh Clinical Academic Track (ECAT) (203913/Z/16/Z). J.P.M and C.J.R are funded by CRUK (A20409, A25142). We thank the CRUK Glasgow Centre and the BSU facilities and Histology Service at the CRUK Beatson Institute. We also thank Scott W. Lowe for pT3-NRas^G12V^-IRES-GFP, pT3-NRas^G12V^/D38A-IRES-GFP and CMVSB13 plasmids. We thank specially Maria Christophorou, Alex von Kriegsheim, Noor Gammoh, Tamir Chandra, Cleo Bishop, Claudia Wellbrock, Nick Hastie, Andrew Jackson, Martin Reijns, Peter Adams, Ana Banito and all the members of J.C.A. lab for helpful criticism, discussion and encouragement.

## SUPPLEMETARY INFORMATION

**Supplemental Figure S1.**
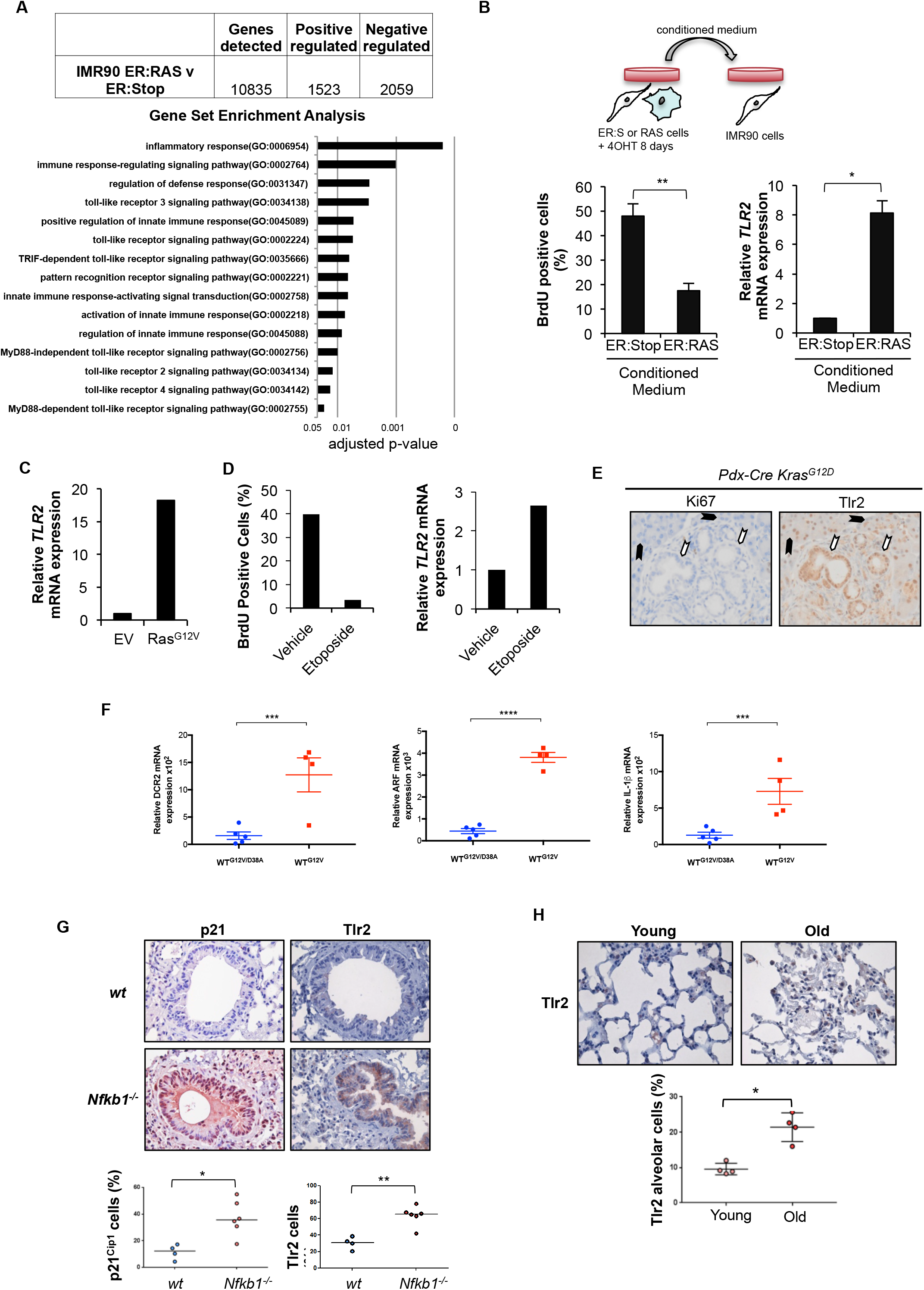
**(A)** Transcriptome analysis and Gene Set Enrichment Analysis was conducted to compare oncogene-induced senescent cells (IMR90 ER:RAS + 4OHT at 8 days) vs. control cells (IMR90 ER:Stop + 4OHT at 8 days) of 6 independent experiments. Adjusted p-value of significant gene sets related to the innate immune response and pattern recognition receptors are shown. **(B)** Conditioned medium (CM) generated by IMR90 ER:Stop and ER:RAS following 8 days of 4OHT treatment was transferred to proliferating IMR90 cells and cultured for 2 days. After a 16-hour pulse, BrdU incorporation was high content analysis. qRT-PCR of *TLR2* mRNA in IMR90 cells treated with CM from ER:RAS and ER:Stop cells was assessed. Results expressed as mean ± SEM of 3 independent experiments. Statistical significance was calculated using Students two-tailed *t*-test. **p < 0.01, * p < 0.05. **(C)** TLR2 mRNA expression determined by qRT-PCR of IMR90 cells infected with RAS^G12V^ vector and empty vector (EV) as control. **(D)** IMR90 cells were treated with 100 µM Etoposide. After 2 days, BrdU incorporation was assessed by immunofluorescence and qRT-PCR analysis for *TLR2* expression was conducted. **(D)** Immunohistochemical staining of PanIN structures from *Pdx-Cre Kras^G12D^* for the expression of Tlr2 and Ki67 expression in consecutive sections. White arrows indicate PanIN structure. Black arrows indicate pancreatic acinar cells. **(F)** qRT-PCR analysis of the senescence markers *dcr2* and *arf* and the key SASP factor *il-1β* from snap frozen liver samples from WT mice 6 days after receiving hydrodynamic delivery of Nras^G12V/D38A^ (n=5) and Nras^G12V^ (n=4) transposons respectively. Scatter plot represents value from individual animals and the horizontal line represents group mean ± SEM. Statistics: Students two-tailed *t*-test. ***p<0.001, ****p<0.0001. **(G)** Analysis of Tlr2 and p21^Cip-1^ expression was conducted by immunohistochemistry in lung sections from wild type (wt) or nfkb1 knock out mice (*nfkb1^-/-^*) at 9.5 months of age. 10-15 random images were captured per mouse and average percentage positivity calculated for airway epithelial compartments. Scatter plots represent mean percentage positivity for each animal with the horizontal line representing group median. Statistical significance was calculated using Mann Whitney U test. *p < 0.05, **p < 0.01. Representative images of p21 and TLR2 staining by immunohistochemistry (positive, brown; negative, blue) in airway epithelial cells from wt and *nfkb1-/-* mice. **(E)** Analysis of Tlr2 expression by immunohistochemistry in lung sections of wt mice at 6.5 months of age (Young) and 24 months of age week (old). Scatter plots were generated from 10-15 random images captured per animal with individual points representing mean percentage positivity for each mouse with horizontal line representing group median. Statistics: Mann-Whitney U test. *p < 0.05. Representative images of TLR2 staining by immunohistochemistry (positive, brown; negative, blue) in alveolar cells from wt mice 6.5 and 24 months of age.

**Supplemental Figure S2.**
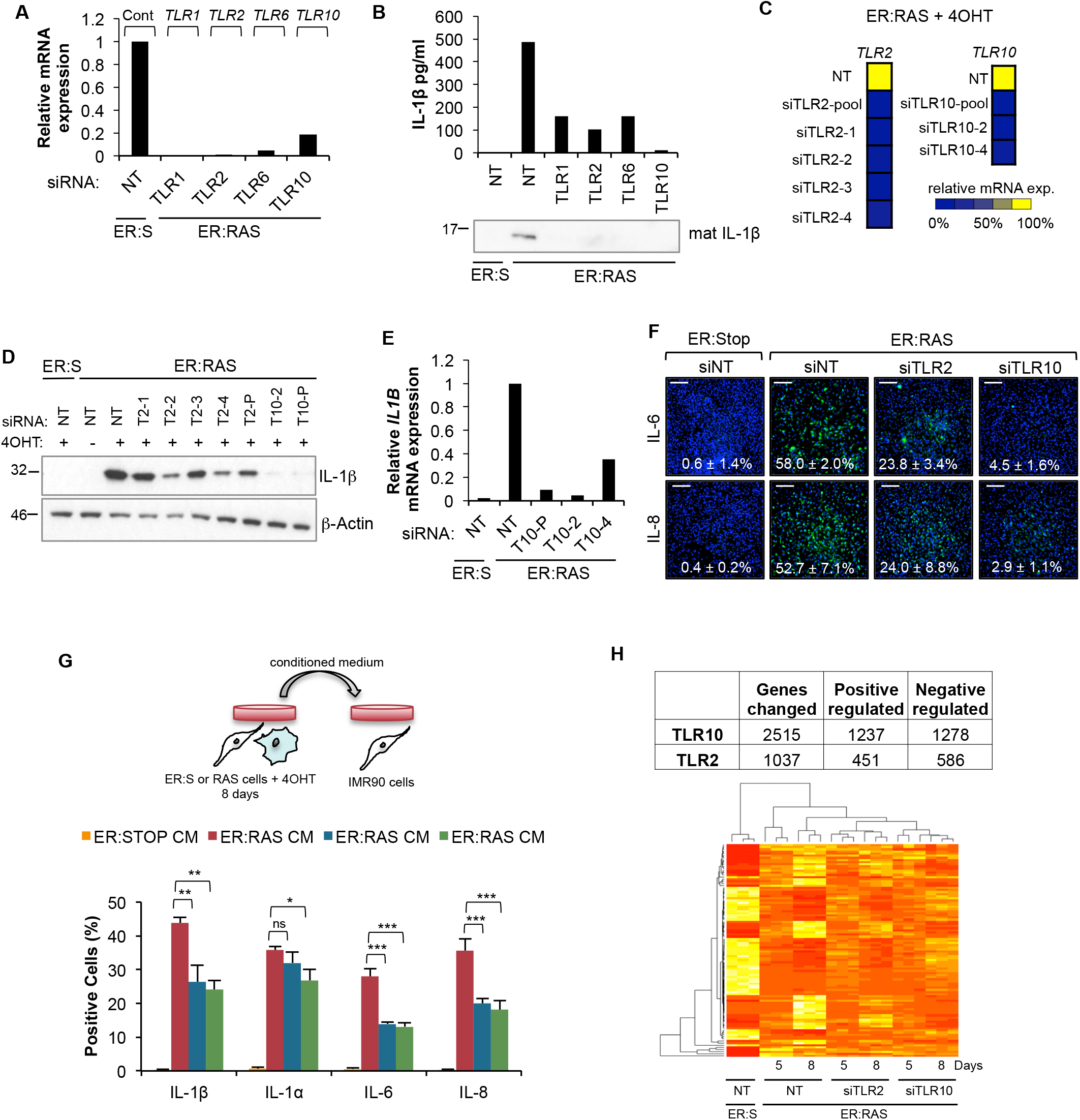
**(A-F)** IMR90 ER:RAS cells were treated with 4OHT and repeatedly transfected with indicated pooled siRNA targeting TLR1 subfamily receptors and non-target (NT) siRNA as control for 8 days. **(A)** qRT-PCR of *TLR1, TLR2, TLR6, TLR10* expression to validate knockdown. **(B)** ELISA and western blot for IL-1β content in the conditioned medium following indicated TLR knockdown. **(C)** qPCR of TLR2 and TLR10 expression following the pooled and individual siRNA media knockdown of respective target genes. **(D)** Western blot of IL-1β protein levels following TLR2 and TLR10 knockdown using pooled (T2-P, T10-P) and individual (T10-2, T2-1, T2-2, T2-3, T2-4) siRNA. **(E)** qRT-PCR of *IL1B* mRNA expression following TLR10 knockdown using pooled (T10-P) and individual (T10-2, T10-4) siRNA. **(F)** Representative images of immunofluorescence and high content analysis of IL6, IL8 and IL-1β expression in TLR2 and TLR10 knockdown cells. **(G)** Conditioned medium (CM) generated by IMR90 ER:Stop and ER:RAS following 8 days of 4OHT treatment was transferred to proliferating IMR90 cells with siRNA knockdown of TLR2 and TLR10 and cultured for 2 days. Immunofluorescence and high content analysis was used to determine IL-1α, IL-1β, IL-6 and IL-8 protein levels in the IMR90 cells. Results expressed as mean ± SEM of 3 independent experiments. Statistical significance was calculated using One-Way ANOVA and Dunnett’s multiple comparison’s tests. ***p < 0.001, **p < 0.01, * p < 0.05, NS, non-significant. **(H)** Transcriptome analysis (Ampliseq) of IMR90 ER:RAS cells transfected with pooled siRNA for TLR2 and TLR10, and non-target (NT) as control. Table identifies the total number of genes significantly regulated by TLR2 and TLR10 during OIS. Heat-map showing sample clustering.

**Supplemental Figure S3.**
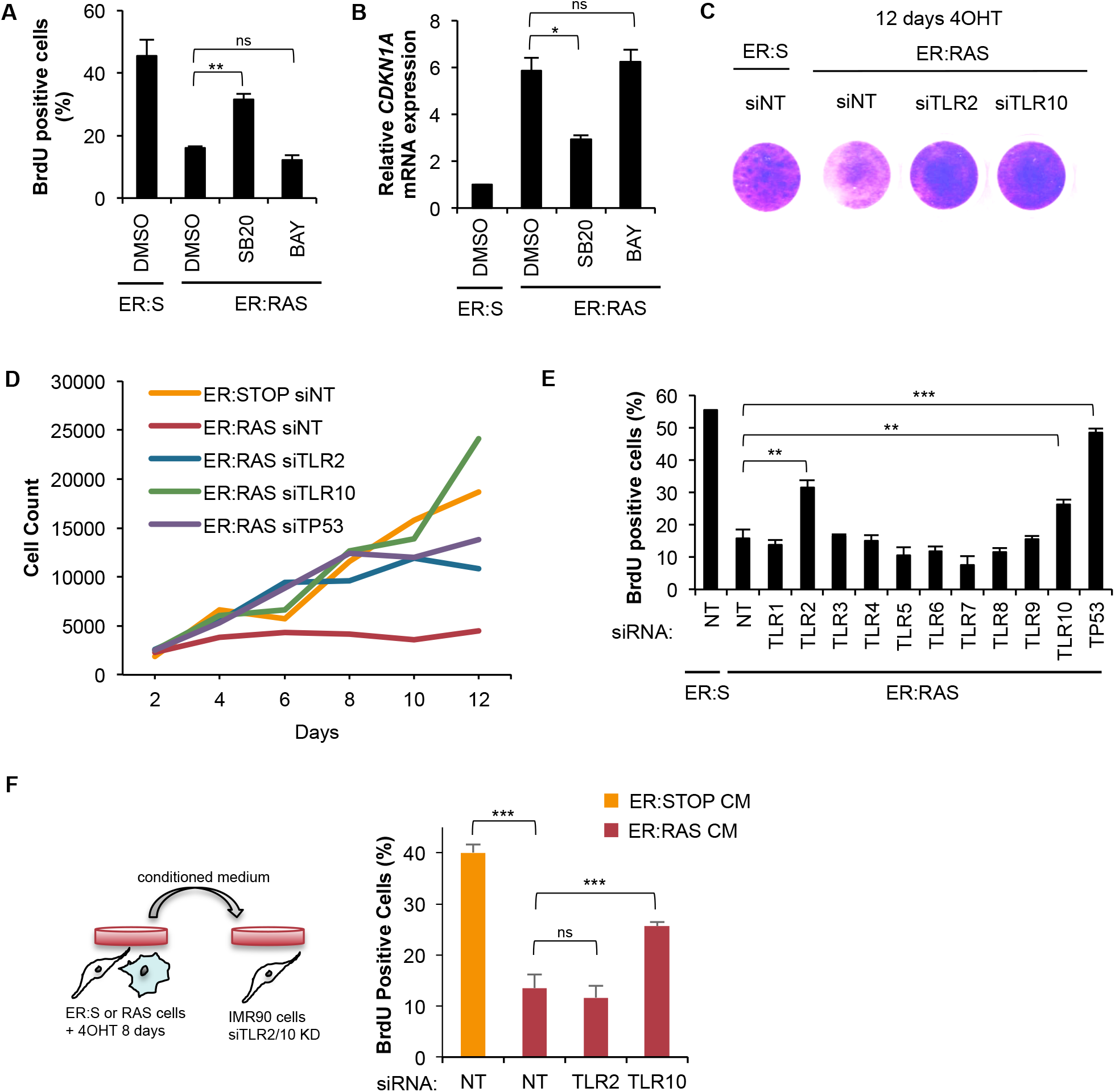
(**A**) IMR90 ER:RAS and ER:Stop cells were treated with 10 μM of IKK inhibitor (BAY 11-07082) and p38 MAPK inhibitor (SB202190). A BrdU incorporation assay was conducted and analysed by high content microscopy. **(B)** *CDKN1A* mRNA expression was measured by qRT-PCR in sample treated as in (A). **(C)** Crystal violet staining of the cells 12 days after siRNA transfection and 4OHT treatment of IMR90 ER:RAS cells. **(D)** Total cell count (nuclei count number) in the total surface of tissue culture wells by high content analysis of IMR90 ER:RAS cells transfected with indicated siRNA for 12 days. **(E)** BrdU incorporation assay following pooled siRNA knockdown of all TLR family members at 5 days of 4OHT treatment in IMR90 ER:RAS and IMR90 ER:Stop cells. Results expressed as mean ± SEM of 3 independent experiments. Statistical significance was calculated using two-tailed Students t-tests. ***p < 0.001, **p < 0.01, * p < 0.05, n.s., non-significant. **(F)** Conditional medium (CM) generated by IMR90 ER:Stop and ER:RAS following 8 days of 4OHT treatment was transferred to proliferating IMR90 cells with siRNA knockdown of TLR2 and TLR10 and cultured for 2 days. Immunofluorescence and high content analysis was used to determine BrdU incorporation in the IMR90 cells. Results expressed as mean ± SEM of 3 independent experiments. Statistical significance was calculated using One-Way ANOVA. ***p < 0.001, **p < 0.01, * p < 0.05, ns, non-significant.

**Supplemental Figure S4.**
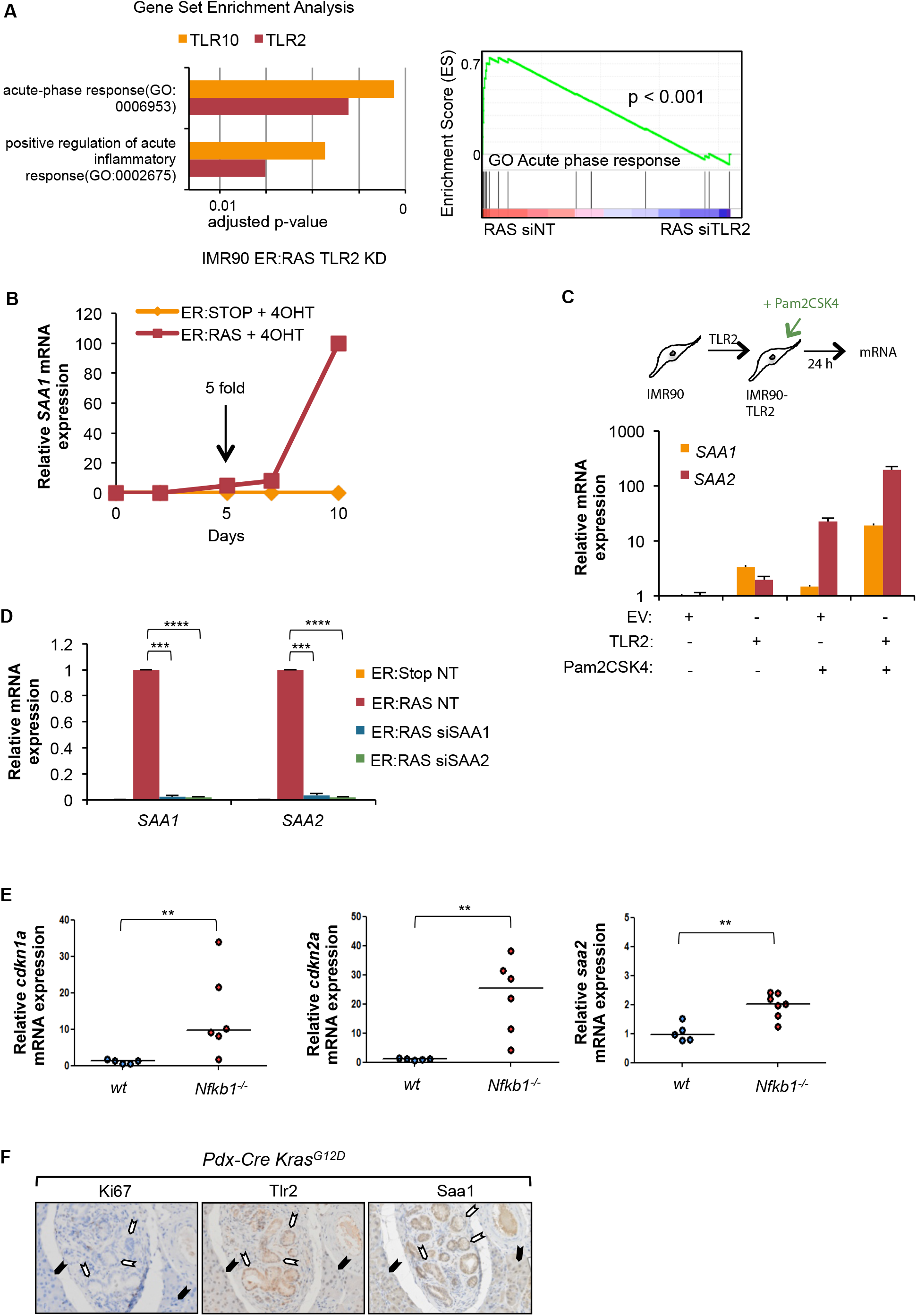
**(A)** Gene Set Enrichment Analysis (GSEA) of the transciptome from Supplemental Fig. S2H showing top two regulated gene set groups, and GSEA enrichment plot of genes of the acute phase response in TLR2 siRNA transfected IMR90 ER:RAS 4OHT induced cells. **(B)** IMR90 ER:RAS and ER:Stop were treated with 4OHT for up to 10 days with RNA collected at indicated intervals. qRT-PCR for SAA1 mRNA expression **(C)** qRT-PCR for SAA1 and SAA2 mRNA expression in TLR2 overexpressing IMR90 cells activated with 1 µg/ml Pam2CSK4 ligand for 3 hours. Results expressed as mean ± SEM of 3 independent experiments. **(D)** Confirmation of *SAA1* and *SAA2* expression knockdown by qRT-PCR in IMR90 ER:RAS and ER:Stop cells transfected with pooled siRNA targeting SAA1 and SAA2, and non-target (NT) control and treated with 4OHT for 8 days. Results expressed as mean ± SEM of 3 independent experiments. All statistical significance was calculated using One-Way ANOVA. ***p < 0.001, **p < 0.01, * p < 0.05, ns, non-significant. **(E)** qRT-PCR for Cdkn1a (p21), Cdkn2a (p16) and Saa2 expression in whole lung tissue from wt or *nfkb1-/-* mice at 9.5 months of age. Dot plots represent the ΔΔCT value for individual animals generated by normalizing to 18S expression. The horizontal line represents group median. Statistical significance was calculated using Mann-Whitney U test. **p < 0.01. **(F)** Immunohistochemical staining of PanIN structures from *Pdx-Cre Kras^G12D^* for the expression of Tlr2, Saa1 and Ki67 expression in consecutive sections. White arrows indicate PanIN structure. Black arrows indicate pancreatic acinar cells.

**Supplemental Figure S5.**
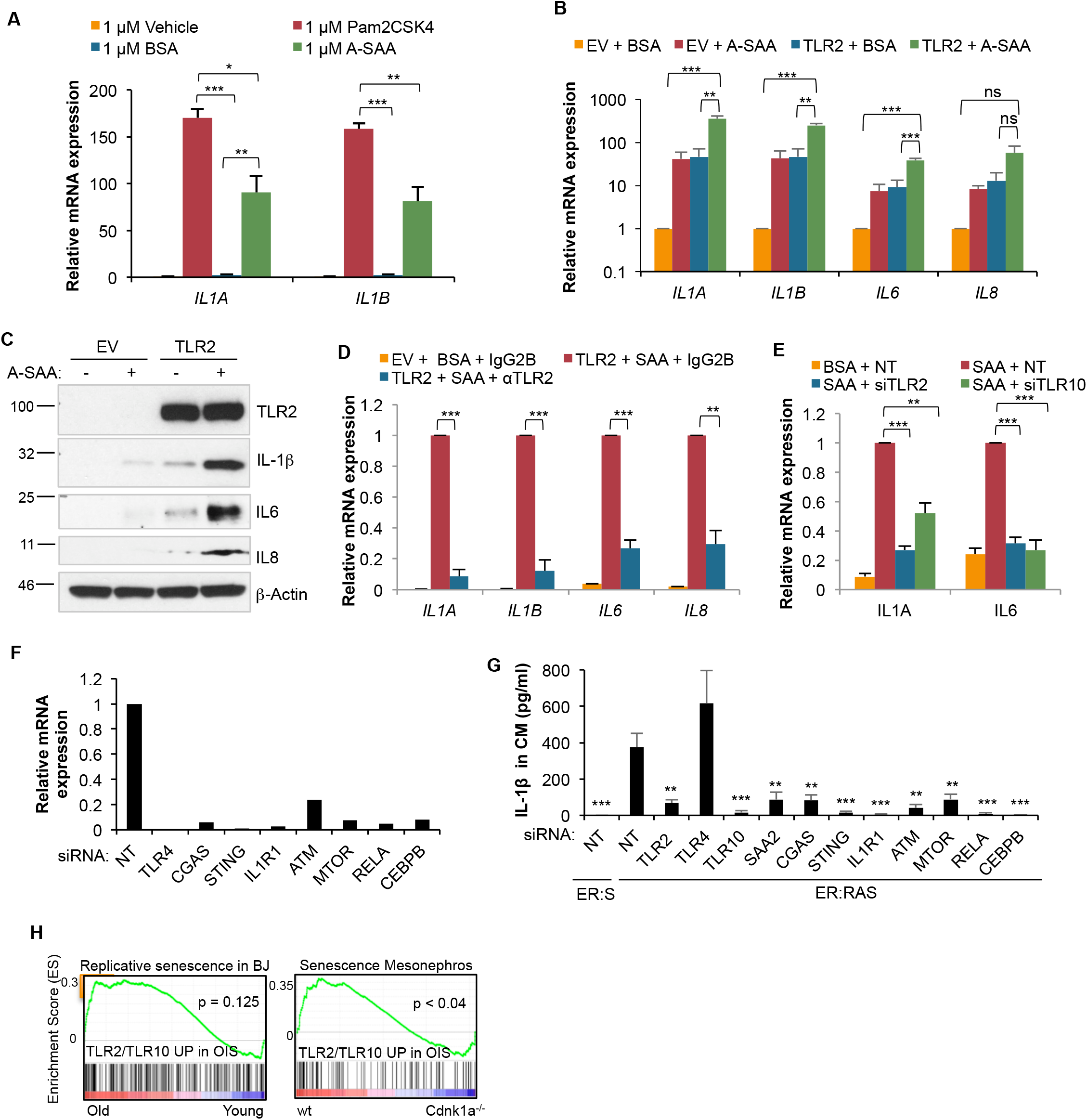
**(A)** *IL1A* and *IL1B* mRNA expression was assessed by qRT-PCR in TLR2 expressing and non-expressing control (EV) IMR90 cells treated with 1 µM A-SAA or 1 µM Pam2CSK4 for 3 hours. Results expressed as mean ± SEM of 3 independent experiments. **(B)** *IL1A, IL1B, IL6* and *IL8* mRNA expression was assessed by qRT-PCR in TLR2 expressing and non-expressing control (EV) IMR90 cells treated with 10 µg/ml A-SAA for 3 hours. Results expressed as mean ± SEM of 3 independent experiments. **(C)** IMR90 cells expressing TLR2 or infected with empty vector (EV) as a control were incubated for 3 hours with recombinant active SAA (A-SAA) protein (10 µg/ml) or BSA (10 µg/ml) as a control. Western blot to measure IL-1β IL6 and IL8 protein levels as indicated. **(D)** TLR2 overexpressing IMR90 cells were treated with 10 µg/ml A-SAA plus 10 µg/ml of neutralising antibody against TLR2 for 2 hours. Assessment of *IL1A, IL1B, IL6* and *IL8* mRNA expression was conducted by qRT-PCR. Results expressed as mean ± SEM of 3 independent experiments. **(E)** IMR90 cells transfected with pooled siRNA for TLR2 and TLR10 were treated with 1 µg/ml A-SAA for 3 hours and *IL6* and *IL1A* mRNA expression determined by qRT-PCR. Results expressed as mean ± SEM of 3 independent experiments. All statistical significance was calculated using One-Way ANOVA. ***p < 0.001, **p < 0.01, * p < 0.05, ns, non-significant. **(F)** qRT-PCR to confirm the knockdown efficiency of the indicated target gene by pooled siRNA. **(G)** IMR90 ER:RAS cells were treated with 4OHT and repeatedly transfected with indicated pooled siRNA and non-target (NT) siRNA as control for 8 days. ELISA for IL-1β levels in conditioned medium. Results expressed as mean ± SEM of 3 independent experiments. **(F)** GSEA plots of the gene set from Supplemental Fig. S4A in the transcriptome of replicative senescence BJ cells and programmed senescent cells of the mesonephros in Fig. 4I.

**Supplemental Figure S6.**
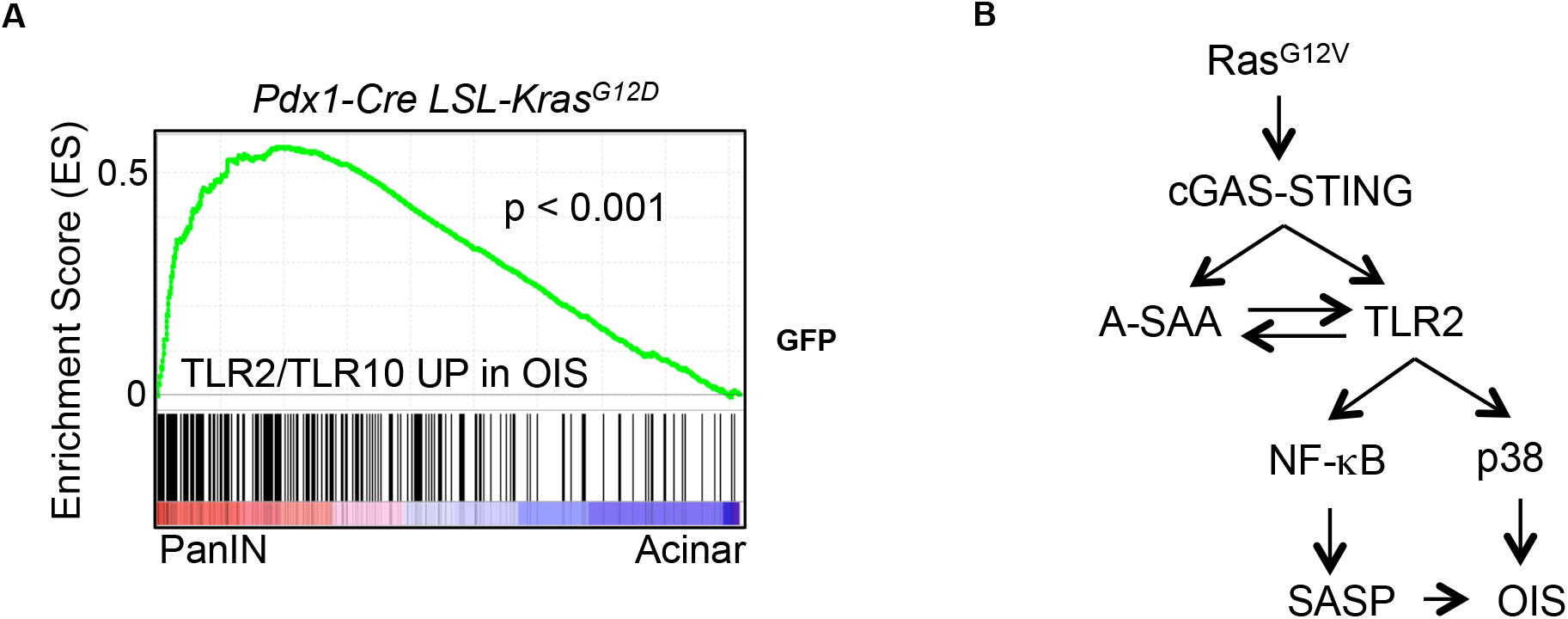
**(A)** GSEA enrichment plot of genes regulated by TLR2 and TLR10 in OIS described in Supplemental figure S2G in the transcriptome of PanIN cells compared to acinar normal cells (GSE33323). **(B)** Model of activation of the SASP by A-SAA-TLR2 in OIS.

**Supplementary table S1.**
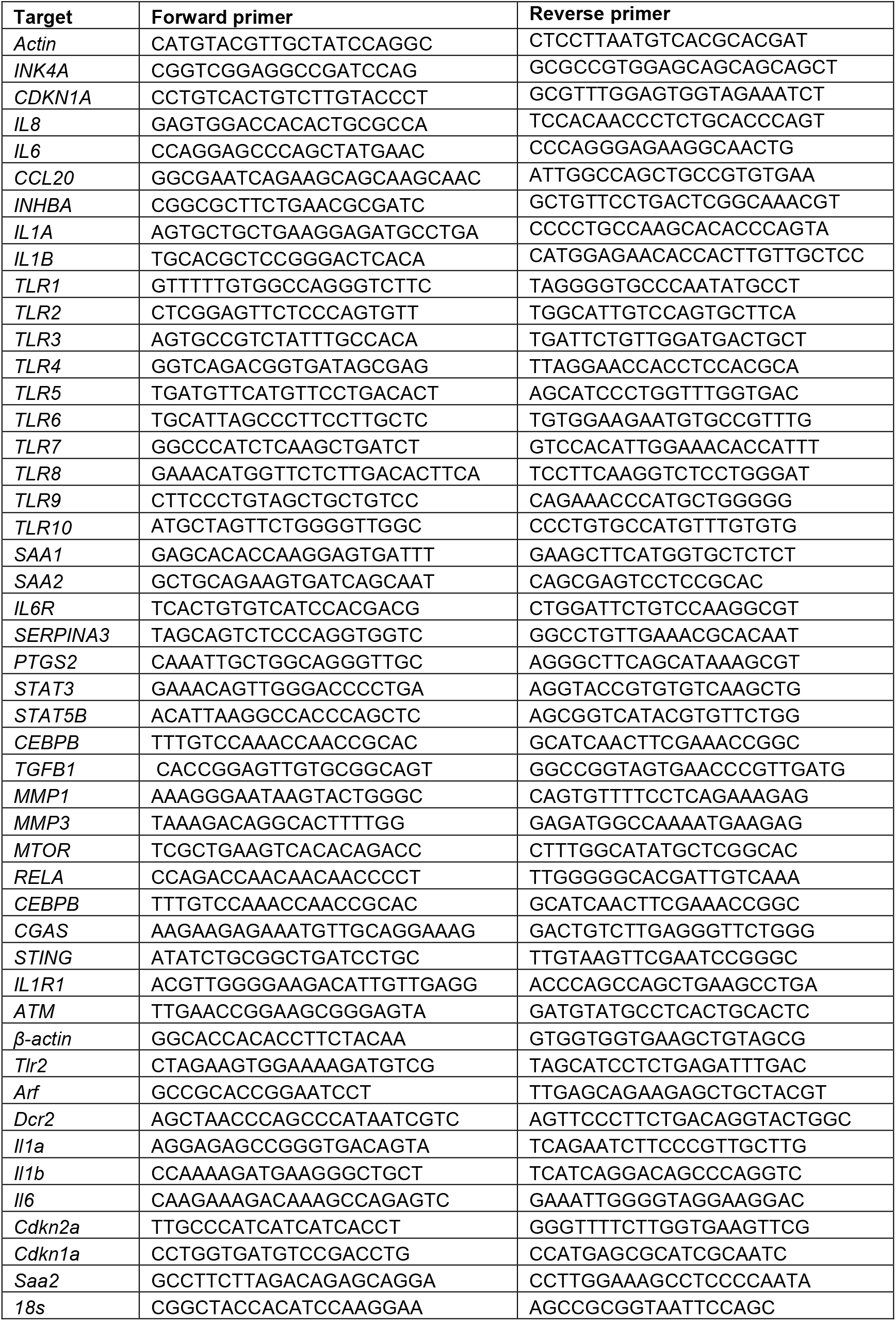
Primers used for RT-qPCR.

**Supplementary table S2.**
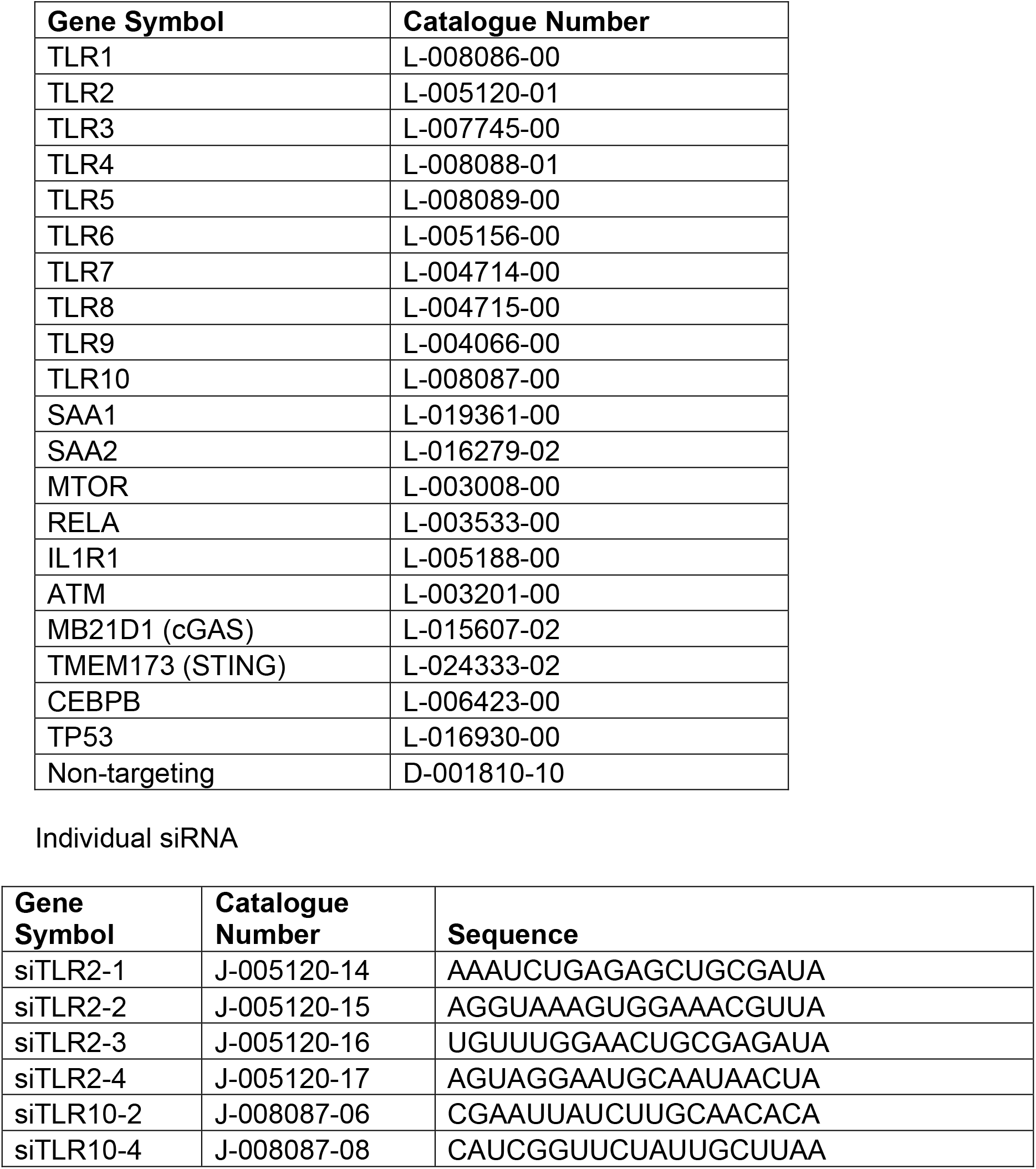
siRNA sequences All ON-TARGETplus siRNA (Dharmacon) SMARTpool.

**Supplementary table S3.** Antibodies.

| REAGENT or RESOURCE | SOURCE | IDENTIFIER | USAGE |
| --- | --- | --- | --- |
| <u>Primary Antibodies</u> |  |  |  |
| Mouse monoclonal anti-IL1A | R&D systems | Cat #MAB200 | WB 1:1000, IF 1:100 |
| Mouse monoclonal anti-IL1B | R&D systems | Cat #MAB201 | WB 1:200 - 1:5000, IF 1:100 |
| Goat polyclonal anti-IL6 | R&D systems | Cat #AF206NA | WB 1:1000, IF 1:100 |
| Mouse monoclonal anti-IL8 | R&D systems | Cat #MAB208 | WB 1:1000, IF 1:100 |
| Mouse monoclonal anti-p53 (DO1) | Santa Cruz | Cat #sc-126 | WB 1:1000, IF 1:100 |
| Rabbit monoclonal anti-TLR2 [EPNCIR133] | Abcam | Cat #ab108998 | WB 1:1000, IF 1:100 |
| Rabbit polyclonal anti-TLR10 | Santa Cruz | Cat #sc-30198 | WB 1:1000 |
| Mouse monoclonal anti-BrdU | BD Phamingene | Cat #555627 | IF 1:2000 |
| Rabbit polyclonal anti-IKKα | CST | Cat #2682 | WB 1:1000 |
| Rabbit polyclonal anti-IKKβ (L570) | CST | Cat #2678 | WB 1:1000 |
| Rabbit monoclonal anti-IKK$\alpha/\beta$ (Ser176/180) (16A6) | CST | Cat #2697 | WB 1:1000 |
| Rabbit monoclonal anti-p38 MAPK (D13E1) XP | CST | Cat #8690 | WB 1:5000 |
| Rabbit polyclonal anti-phospho-p38 MAPK (Thr180/Tyr182) | CST | Cat #9211 | WB 1:1000 |
| Rabbit monoclonal anti-STING (D2P2F) | CST | Cat #13647 | WB 1:1000 |
| Rabbit monoclonal anti-phospho NF-kB p65 (Ser536) | CST | Cat #3033 | WB 1:500 |
| Mouse monoclonal anti-NF-kB p65 (F-6) | Santa Cruz | Cat #sc-8008 | WB 1:1000 |
| Mouse monoclonal anti-TLR2 | R&D systems | Cat #MAB2616 | Neut 10 μg/ml |
| Mouse monoclonal IgG2B Isotype Controls (20116) | R&D systems | Cat #MAB004 | Neut 10 μg/ml |
| Rabbit monoclonal anti-Ki67 [SP6] | Abcam | Cat #ab21700 | IHC |
| Rabbit polyclonal anti-TLR2 | Invitrogen | Cat #PA5-20020 | IHC 1:400 |
| Rabbit polyclonal anti-SAA1 | Biorbyt | Cat #orb228668 | IHC 1:200 |
| Rabbit polyclonal anti-p21 | Abcam | Cat #ab7960 | IHC 1:200 |
| Mouse monoclonal anti-N-Ras (F155) | Santa Cruz | Cat #sc-31 | IHC 1:300 |
| Rabbit polyclonal anti-IL-1b (H-153) | Santa Cruz | Cat #sc-7884 | IHC 1:100 |
| Rabbit polyclonal anti-GFP | Abcam | Cat#ab6556 | IHC 1:1000 |
| Goat polyclonal anti-IL1A | R&D systems | Cat#AF-400 | IHC 1:200 |
| <u>Secondary Antibodies</u> |  |  |  |
| Goat anti-Mouse IgG Alexa Fluor 488 | Thermo Fisher | Cat #A11029 | IF 1:1000 |
| Donkey anti-Goat IgG Alexa Fluor 594 | Thermo Fisher | Cat #A-11058 | IF 1:1000 |
| Anti-Mouse IgG (Fc specific)-Peroxidase | Sigma | Cat #A2554 | WB 1:20,000 - 1:80,000 |
| Anti-Rabbit IgG (whole molecule)-Peroxidase | Sigma | Cat #A0545 | WB 1:20,000 - 1:80,000 |
| Donkey anti-goat-HRP | Santa Cruz | Cat #sc-2020 | WB 1:20,000 |
| Goat anti-Rabbit IgG Alexa Fluor 594 | Thermo Fisher | Cat #A11037 | IF 1:1000 |
| Monoclonal anti-B-Actin-peroxidase (AC-15) | Sigma | Cat #A3854 | WB 1:100,000 - 1:1,000,000 |

